# Homeostatic macrophages prevent preterm birth and improve neonatal outcomes by mitigating *in utero* sterile inflammation

**DOI:** 10.1101/2023.07.19.549717

**Authors:** Valeria Garcia-Flores, Zhenjie Liu, Roberto Romero, Roger Pique-Regi, Yi Xu, Derek Miller, Dustyn Levenson, Jose Galaz, Andrew D. Winters, Marcelo Farias-Jofre, Jonathan J. Panzer, Kevin R. Theis, Nardhy Gomez-Lopez

## Abstract

Preterm birth (PTB), often preceded by preterm labor, is a major cause of neonatal morbidity and mortality worldwide. Most PTB cases involve intra-amniotic inflammation without detectable microorganisms, termed *in utero* sterile inflammation, for which there is no established treatment. Here, we propose homeostatic macrophages to prevent PTB and adverse neonatal outcomes caused by *in utero* sterile inflammation. Single-cell atlases of the maternal-fetal interface revealed that homeostatic maternal macrophages are reduced with human labor. M2 macrophage treatment prevented PTB and reduced adverse neonatal outcomes in mice with *in utero* sterile inflammation. Specifically, M2 macrophages halted premature labor by suppressing inflammatory responses in the amniotic cavity, including inflammasome activation, and mitigated placental and offspring lung inflammation. Moreover, M2 macrophages restored neonatal gut homeostasis and enhanced resistance to systemic bacterial infection. Our findings show that M2 macrophages are a promising strategy to mitigate PTB and improve neonatal outcomes from *in utero* sterile inflammation.

## INTRODUCTION

Preterm birth, the leading cause of neonatal morbidity and mortality worldwide^1,2^, is often preceded by spontaneous preterm labor, a syndrome of multiple etiologies^3^. Among the known and proposed causes of preterm labor, intra-amniotic inflammation is the best characterized and accounts for a large proportion of cases^3–5^. The recent incorporation of next generation sequencing in obstetrics has revealed that most cases of intra-amniotic inflammation occur in the absence of invading microbes in the amniotic cavity^6–9^, resulting in the discovery of sterile intra-amniotic inflammation (hereafter referred to as *in utero* sterile inflammation). Hence, this new condition is diagnosed by elevated concentrations of inflammatory mediators such as interleukin (IL)-6 in amniotic fluid, in the absence of detectable microorganisms using culture and molecular microbiological techniques^6–10^. Importantly, *in utero* sterile inflammation has been linked to adverse short- and long-term outcomes for the offspring of women with this clinical condition^7,11^. Specifically, women with *in utero* sterile inflammation are at greater risk of having placentas affected by acute histologic chorioamnionitis^12^, which is linked to the development of deleterious neonatal conditions such as bronchopulmonary dysplasia^13–15^ and necrotizing enterocolitis^16,17^, likely due to exposure of the fetal lungs and intestine to intra-amniotic inflammation^18–20^. However, despite the strong associations between *in utero* sterile inflammation and adverse fetal and neonatal outcomes, no approved treatments currently exist.

Sterile inflammation can be triggered by danger signals or alarmins released during cellular stress or injury^21–23^. Consistently, clinical studies have also shown that women with spontaneous preterm labor and *in utero* sterile inflammation have increased amniotic fluid concentrations of alarmins^7,24–26^. Indeed, women with preterm labor and elevated amniotic concentrations of the prototypical alarmin high-mobility group box-1 (HMGB1) (≥8.55 ng/mL) delivered earlier than those with lower concentrations of this alarmin^7^. Furthermore, we have provided mechanistic evidence showing that the *in utero* delivery of HMGB1 induces preterm labor and birth in mice^27–29^. Moreover, our *in vitro* studies demonstrated that incubation with HMGB1 induces the activation of the NLRP3 inflammasome^30^, one of the central pathway in triggering preterm labor and birth in women^31^ and mice^32–34^ experiencing *in utero* sterile inflammation^35^. Hence, a therapeutic approach targeting the NLRP3 inflammasome, elevated inflammatory cytokines in amniotic fluid such as IL-6, and the process of preterm labor could represent a promising strategy for treating *in utero* inflammation and its devastating perinatal consequences.

The maternal-fetal interface hosts a notable and heterogeneous population of macrophages^36–43^. Specifically, we reported that macrophages expressing a homeostatic or M2-like phenotype are more abundant in both term and preterm gestation than those expressing inflammatory phenotypes^42^, pointing to an important role for these cells in maintaining pregnancy homeostasis. Next, we demonstrated that the depletion of maternal macrophages results in preterm birth as well as neonatal growth restriction and increased mortality^43^. Furthermore, we also showed that the adoptive transfer of M2-polarized macrophages prevents preterm birth induced by intra-amniotic LPS^43^, providing proof-of-concept that such cells can serve as a therapeutic approach for *in utero* sterile inflammation. Thus, here we propose the use of M2-polarized macrophages as a cellular therapy to prevent preterm labor associated with *in utero* sterile inflammation as well as its consequences for the offspring.

In this study, we employ a translational mechanistic approach by first leveraging our single-cell atlases of the human maternal-fetal interface to demonstrate a labor-associated reduction of homeostatic macrophages. Next, by using a clinically relevant animal model of *in utero* sterile inflammation induced by the intra-amniotic injection of the alarmin HMGB1, we investigate the potential restoration of M2-polarized macrophages (hereafter also referred to as M2 macrophages) via adoptive transfer to prevent preterm birth and reduce the adverse neonatal outcomes. In addition, we utilize molecular approaches to investigate the inflammatory responses driven by HMGB1-induced *in utero* sterile inflammation and the homeostatic effects of M2 macrophage treatment in maternal and fetal tissues targeted by transferred macrophages, including those involved in the common pathway of parturition. Moreover, we evaluate the damage to key fetal and neonatal organs, namely the lung and intestine, driven by exposure to *in utero* sterile inflammation, including alterations of the gut microbiome, and whether this was reverted by M2 macrophage treatment. Last, we challenge neonates with Group B *Streptococcus* to determine whether M2 macrophage treatment restores neonatal immunocompetence. Collectively, our data indicate that treatment with M2 macrophages represents a novel cellular approach that can prevent preterm birth and ameliorate the adverse neonatal outcomes induced by *in utero* sterile inflammation.

## RESULTS

### The maternal-fetal interface hosts a homeostatic macrophage population that is diminished with labor

We first hypothesized that labor is accompanied by a reduction of homeostatic macrophages in maternal compartments. Our previous flow cytometry studies targeting specific macrophage subsets suggested that such a reduction occurs at the maternal-fetal interface (i.e., the decidua)^42,43^. However, we sought to test our hypothesis using an unbiased discovery approach. To do this, we leveraged our previously generated single-cell atlases of the myometrium and maternal-fetal interface^44–46^. The maternal-fetal interface includes key sites of contact between maternal and fetal tissues: the fetal placenta embedded in the maternal decidua basalis adjacent to the myometrium, and the fetal extraplacental membranes enclosing the amniotic cavity and attached to the maternal decidua parietalis, next to the myometrium (Figure 1A). Our myometrial single-cell atlas includes samples collected from women with term labor as well as term non-labor controls^45^, whereas our single-cell atlases of the placenta and extraplacental membranes also include samples from women with preterm labor in addition to the term groups^44,46^. Given that preterm non-labor deliveries are only performed due to pregnancy complications, such cases are not suited for use as gestational age controls for preterm labor and thus historically have not been utilized in our studies. After normalizing data from our three single-cell atlases, we identified seven distinct macrophage clusters, termed M1 – M7, across all tissues (Figure 1B, Supplementary Table 1). Of note, these cluster numbers were not chosen to correlate with the conventional M1-M2 paradigm, but rather reflect cluster number assignments. We focused on the M1 and M2 clusters because they constituted a significant proportion of macrophages in the myometrium, placenta, and extraplacental membranes (Figure 1B).

**Figure 1.**
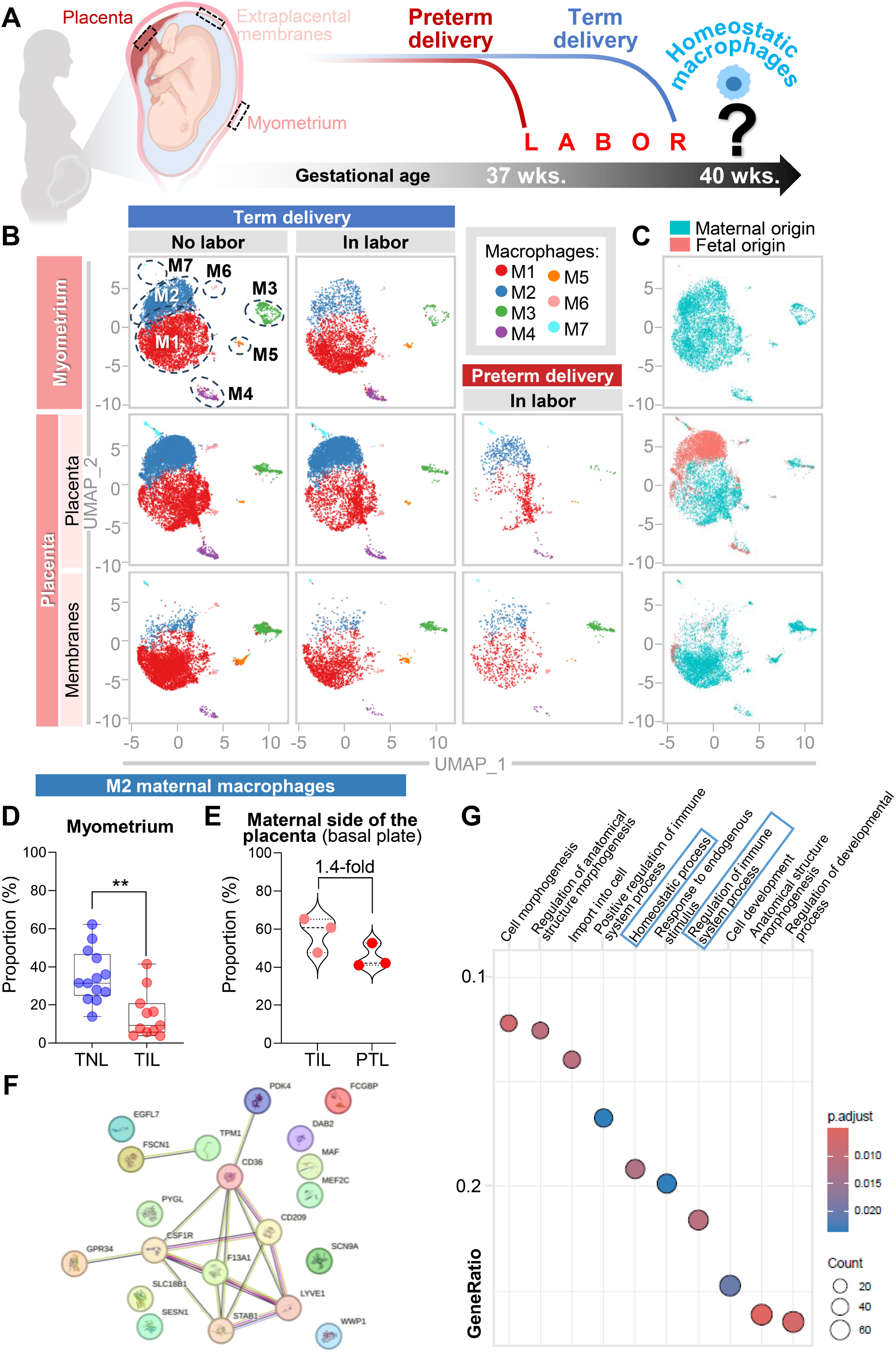
Single-cell atlases of the human maternal-fetal interface reveal that maternal homeostatic macrophages are reduced with labor. **(A)** Representative diagram showing the placenta, extraplacental membranes, and myometrium, along with our working hypothesis that a reduction in homeostatic macrophages may accompany the processes of labor. **(B)** Uniform Manifold Approximation and Projection (UMAP) plots showing seven distinct macrophage clusters (M1 – M7) in (*Top row*) the myometrium from women who delivered at term with (n=11) or without (n=13) labor, (*Middle row*) the placenta (placental villi and basal plate) from women who delivered at term with (n=27) or without (n=21) labor or preterm after preterm labor (n=3), and (*Bottom row*) extraplacental membranes from women who delivered at term with (n=27) or without (n=21) labor or preterm after preterm labor (n=3). **(C)** UMAP plots showing maternal (blue) or fetal (red) origin of all macrophage clusters in the (*top to bottom*) myometrium, placenta, and extraplacental membranes. **(D)** Plot showing the proportions of maternal M2 macrophages in the myometrium from women with term non-labor (TNL) or those with term labor (TIL). **(E)** Plot showing the proportions of maternal M2 macrophages in the basal plate from women with term in labor (TIL) or preterm labor (PTL). P-values were determined using Mann-Whitney U-tests. **p < 0.01. (F) STRING analysis showing the top 20 marker genes from the M2 cluster. (G) Over-representation analysis showing enriched Gene Ontology processes in the M2 cluster. Dot size corresponds to gene count and color scaling represents false discovery rate-adjusted p-values (q<0.05) as determined by Wilcoxon Rank Sum test. See also Figure S1 and Table S1.

Our single-cell atlases include maternal and fetal genotyping data, which allow us to assign an origin to each individual cell. Therefore, we next determined the origins of macrophages in each compartment (Figure 1C). As expected, macrophages in the myometrium were entirely of maternal origin (Figure 1C). In the placenta, both maternal and fetal macrophages were identified, with the majority of maternal macrophages corresponding to the M1 cluster and fetal macrophages to the M2 cluster (Figure 1C). By contrast, the extraplacental membranes include some fetal macrophages, but the M2 cluster was predominantly of maternal origin (Figure 1C). Next, we evaluated whether the proportions of M1 and M2 clusters, as well as other macrophage clusters, differed between labor and non-labor samples. The only subset that decreased during labor was the maternal M2 cluster in the myometrium (Figure 1D). We also observed a non-significant 1.4-fold decrease in the proportion of the maternal M2 cluster in the basal plate (maternal tissue attached to the placenta) in women with preterm labor and birth compared to those with term labor and birth (Figure 1E). This may be due to the small sample size and the challenges associated with collecting human preterm samples. Last, using the top 20 marker genes for the M2 cluster (Figure 1F), we performed over-representation analysis (ORA) based on the Gene Ontology (GO). The biological processes enriched in the M2 cluster included "homeostatic process," "regulation of immune system process," and "regulation of developmental process," supporting the homeostatic functions of this macrophage cluster (Figure 1G and Supplementary Figure 1A). These processes were distinct from those enriched in other macrophage clusters (Supplementary Figure 1B).

Taken together, our single-cell atlases confirmed that the myometrium and other maternal compartments attached to the placenta host a significant population of homeostatic macrophages, and that these cells are reduced during labor, confirming our hypothesis (Figure 1A).

### M2 macrophage treatment prevents preterm birth and improves neonatal survival in a mouse model of *in utero* sterile inflammation

Next, we determined whether homeostatic macrophages could serve as a viable therapeutic approach to prevent preterm birth. We therefore performed adoptive transfer of M2 macrophages in a murine model of *in utero* sterile inflammation induced by the ultrasound-guided intra-amniotic injection of the alarmin HMGB1 (Figure 2A). To ensure the clinical relevance of this model, we injected HMGB1 at concentrations found in women with preterm labor and *in utero* sterile inflammation^7^. The intra-amniotic delivery of HMGB1 shortened gestational length (Figure 2B), resulting in high rates of preterm birth (Figure 2C). Notably, treatment with M2 macrophages extended gestational length, preventing preterm birth (Figure 2B&C). Neonatal mortality was elevated in mice exposed to *in utero* sterile inflammation, given that most preterm neonates die; yet, such mortality was mitigated by treatment with M2 macrophages (Figure 2D). Furthermore, neonates born to dams intra-amniotically injected with HMGB1 failed to thrive, but once again this effect was ameliorated by treatment with M2 macrophages (Figure 2E). Moreover, surviving neonates that had been exposed to HMGB1-induced *in utero* sterile inflammation displayed growth restriction; however, such impairment was rescued by prenatal treatment with M2 macrophages (Figure 2F). Thus, these data indicate that M2 macrophage treatment can serve as a viable therapy to not only prevent preterm birth but also ameliorate the adverse neonatal outcomes driven by exposure to *in utero* sterile inflammation.

**Figure 2.**
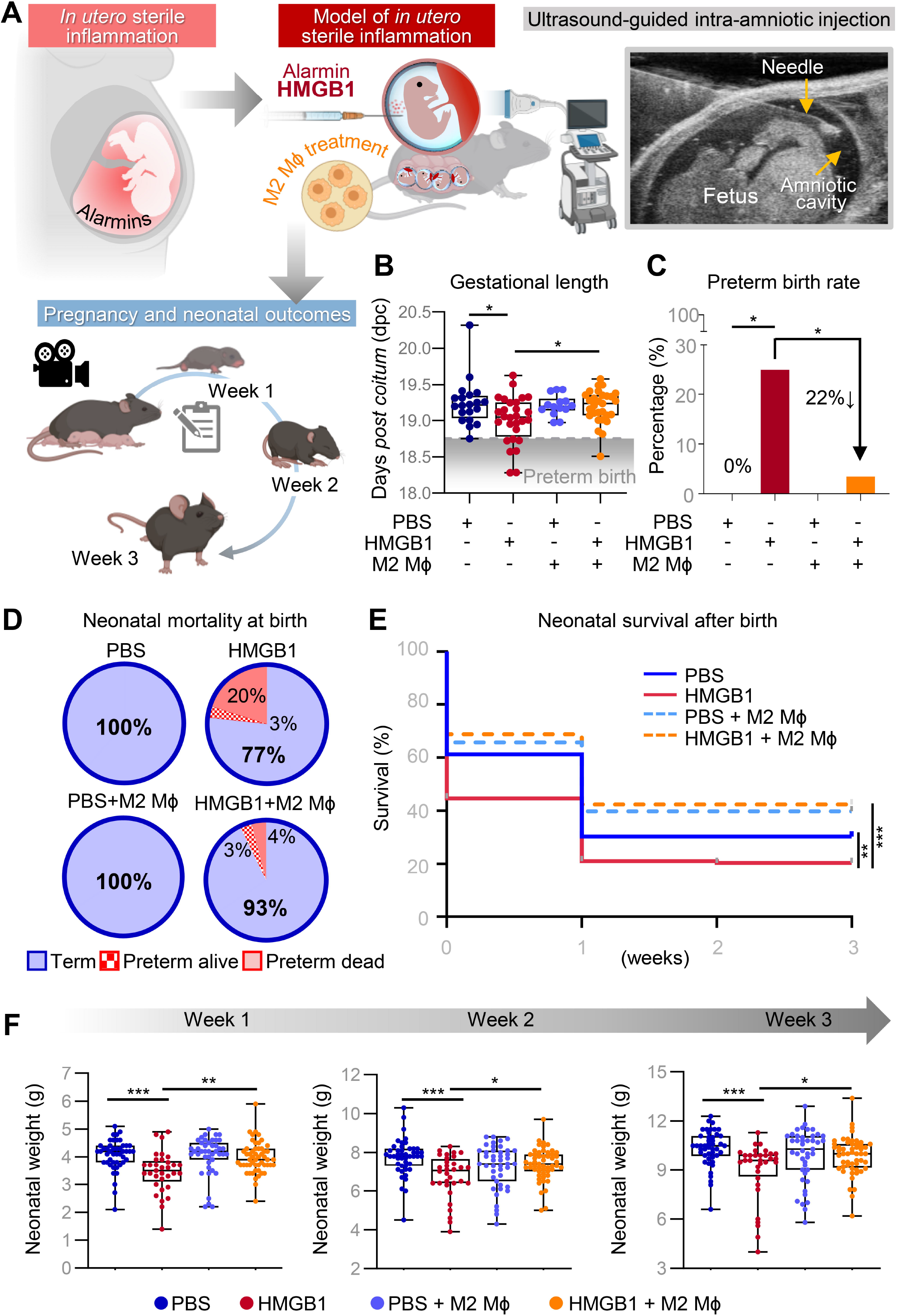
M2 macrophages prevent preterm birth and neonatal mortality induced by *in utero* sterile inflammation. **(A)** To induce *in utero* sterile inflammation, the alarmin HMGB1 was intra-amniotically administered to C57BL/6 dams under ultrasound guidance on 14.5 days *post coitum* (dpc). Bone marrow-derived cells were collected from C57BL/6 mice, differentiated, and polarized to an M2 phenotype (M2 MΦ) *in vitro*. M2 MΦ were administered intravenously to C57BL/6 dams on 13.5 and 14.5 dpc. Dams were monitored until delivery, and neonatal survival and weight were recorded until three weeks of age. **(B)** Gestational length shown as box plots where midlines indicate medians and whiskers indicate minimum/maximum range. P-values were determined using the Kruskal-Wallis test followed by two-stage linear step-up procedure of Benjamini, Krieger, and Yekutieli post-hoc test. **(C)** Rates of preterm birth among dams injected with PBS (n = 20), HMGB1 (n = 28), PBS + M2 MΦ (n = 14), and HMGB1 + M2 MΦ (n = 29) are shown as bar plots. P-values were determined using two-sided Fisher’s exact test. **(D)** Pie charts representing the survival at birth of preterm neonates. PBS (n = 20 litters), HMGB1 (n = 20 litters), PBS + M2 MΦ (n = 14 litters), and HMGB1 + M2 MΦ (n = 16 litters). **(E)** Kaplan-Meier survival curves from neonates at weeks 1, 2, and 3 of life. PBS (n = 20 litters), HMGB1 (n = 20 litters), PBS + M2 MΦ (n = 14 litters), and HMGB1 + M2 MΦ (n = 16 litters). P-values were determined using the Gehan-Breslow-Wilcoxon test. **(F)** Individual weights (grams, g) of neonates across the first three weeks of life are shown as box plots where midlines indicate medians and whiskers indicate minimum/maximum range. PBS (n = 11 litters), HMGB1 (n = 11 litters), PBS + M2 MΦ (n = 11 litters), and HMGB1 + M2 MΦ (n = 14 litters). P-values were determined using the Kruskal-Wallis test followed by Dunn’s post-hoc test. *p < 0.05; **p < 0.01; ***p < 0.001.

### M2 macrophages dampen HMGB1-induced *in utero* sterile inflammation, including inflammasome activation, in the amniotic cavity

We next sought to uncover the mechanisms whereby adoptively transferred M2 macrophages were preventing adverse outcomes driven by HMGB1-induced *in utero* sterile inflammation. First, to establish the kinetics of the *in utero* sterile inflammatory response, we collected the amniotic fluid of HMGB1-injected dams at 24, 48, 72, or 96 h post-injection to evaluate the concentrations of IL-6, total IL-1β and TNF - classic cytokines associated with *in utero* sterile inflammation^25,26^ (Figure 3A). The intra-amniotic delivery of HMGB1 increased the amniotic fluid concentrations of IL-6, the gold standard cytokine used to clinically diagnose *in utero* sterile inflammation^6,7^, at 72 h post-injection (Figure 3B), consistent with the observed timing of preterm labor post-HMGB1 injection. Notably, M2 macrophage treatment not only reduced amniotic fluid concentrations of IL-6 at 72 h post-injection, but also those of TNF and total IL-1β (Figure 3C&D), even though these were not significantly affected by HMGB1 (Supplementary Figure 2). We then focused on the inflammasome, a key signaling pathway implicated in the *in utero* inflammatory response triggered by alarmins, leading to the processing of active caspase (CASP)-1 and mature IL-1β (Figure 3E)^31–33,35^. The intra-amniotic delivery of HMGB1 caused an increase in active CASP-1 and mature IL-1β in amniotic fluid (Figure 3F&G). Notably, M2 macrophage treatment dampened such inflammasome activation by reducing both active CASP-1 and mature IL-1β (Figure 3H&I). These data indicate that M2 macrophages exert their homeostatic effects in the amniotic cavity by dampening the HMGB1-induced inflammatory cytokine response, including inflammasome activation.

**Figure 3.**
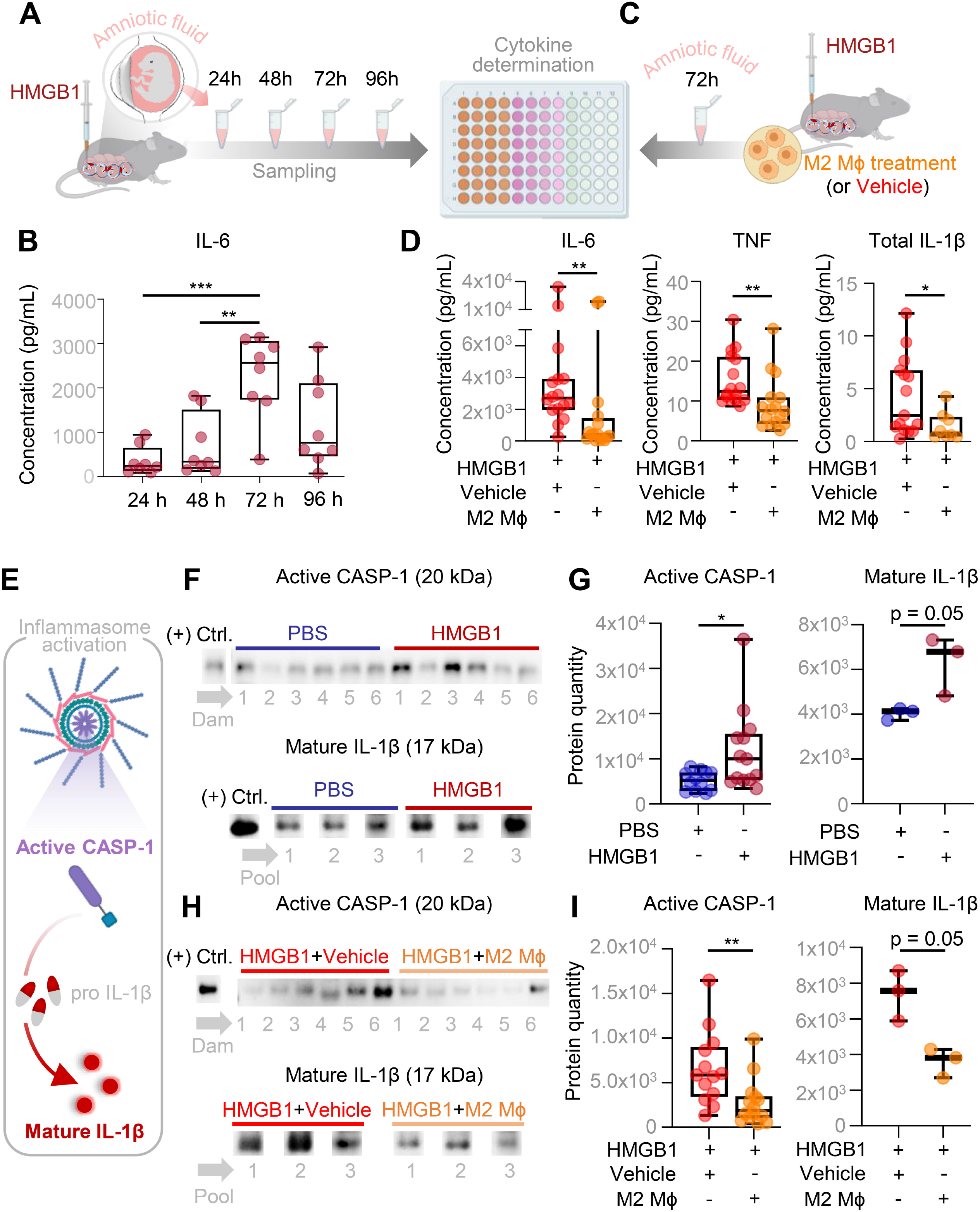
M2 macrophages dampen HMGB1-induced *in utero* sterile inflammation, including inflammasome activation, in the amniotic cavity. **(A)** Dams were intra-amniotically injected with HMGB1 on 14.5 days *post coitum* (dpc). Amniotic fluid was collected at 24, 48, 72, or 96 h post-HMGB1 injection for cytokine determination. **(B)** Concentrations of IL-6 in the amniotic fluid of HMGB1-injected dams at 24, 48, 72, and 96 h post-injection are shown as box plots (n = 8 dams per time point). P-values were determined using the Kruskal-Wallis test followed by two-stage linear step-up procedure of Benjamini, Krieger, and Yekutieli post-hoc test. **(C)** M2 macrophages (M2 MΦ) or vehicle (PBS) were intravenously administered on 13.5 dpc and 14.5 dpc to dams followed by ultrasound-guided intra-amniotic injection of HMGB1 on 14.5 dpc. Amniotic fluid was collected at 72 h post-HMGB1 injection for cytokine determination. **(D)** Concentrations of IL-6, TNF, and total IL-1β in the amniotic fluid of the HMGB1+Vehicle dams (n = 16 dams) or HMGB1+M2 MΦ dams (n = 16 dams) at 72 h post-injection are shown as box plots. P-values were determined using the two-tailed Mann-Whitney U-test. **(E)** To determine inflammasome activation, amniotic fluid of dams that received PBS, HMGB1, HMGB1 + Vehicle, or HMGB1 + M2 MΦ was collected at 72 h post-HMGB1 injection for immunoblotting. **(F)** Immunoblotting of active caspase (CASP)-1 expression and mature IL-1β expression in the amniotic fluid of dams injected with PBS or HMGB1. Representative CASP-1 immunoblot image shows 6 samples per group, and representative mature IL-1β immunoblot image shows 3 pooled samples (pooled amniotic fluids from 3 dams) per group. **(G)** Protein quantification of active CASP-1 in the amniotic fluid of dams injected with PBS (n = 12) or HMGB1 (n = 13). Protein quantification of mature IL-1β in the pooled amniotic fluid of dams injected with PBS (n = 3) or HMGB1 (n = 3). Data are shown as boxplots where midlines indicate medians, boxes denote interquartile ranges, and whiskers indicate the minimum/maximum range. P-values were determined using the one-tailed Mann-Whitney U-test. **(H)** Immunoblotting of active CASP-1 expression and mature IL-1β expression in the amniotic fluid of HMGB1+Vehicle or HMGB1+M2 MΦ dams. Representative CASP-1 immunoblot image shows 6 samples per group, and representative mature IL-1β immunoblot image shows 3 pooled samples (pooled amniotic fluids from 3 dams) per group. **(I)** Protein quantification of active CASP-1 in the amniotic fluid of dams injected with HMGB1+Vehicle (n = 12) or HMGB1+M2 MΦ (n = 14). Protein quantification of mature IL-1β in the pooled amniotic fluid of dams injected with HMGB1+Vehicle (n = 3) or HMGB1+M2 MΦ (n = 3). P-values were determined using the one-tailed Mann-Whitney U-test. Data are shown as boxplots where midlines indicate medians, boxes denote interquartile ranges, and whiskers indicate the minimum/maximum range. *p < 0.05; **p < 0.01; ***p < 0.001. See also Figure S3.

### M2 macrophages interfere with the pathway of preterm labor

The processes of term and preterm labor share a common underlying pathway including fetal membrane activation, uterine contractility, and cervical remodeling^47–50^. Such processes are characterized by the activation of pro-inflammatory signaling networks, including inflammasome activation^51^. Indeed, we recently reported that inflammasome activation in the fetal membranes and uterine tissues is essential for the onset of preterm labor in mice undergoing *in utero* sterile inflammation^33^. However, this pathway is not involved in the cervical processes associated with preterm labor^33,52^. Hence, we next evaluated whether M2 macrophage treatment abrogates inflammatory processes, including inflammasome activation in the fetal membranes (Figure 4A) and uterine tissues. Consistent with our findings in amniotic fluid, intra-amniotic injection of HMGB1 increased levels of active CASP-1 and mature IL-1β in the fetal membranes, indicating inflammasome activation (Figure 4B&C). Yet, M2 macrophage treatment abrogated inflammasome activation in the fetal membranes, as indicated by reduced quantities of active CASP-1 and mature IL-1β (Figure 4D&E). Moreover, gene expression profiling revealed that intra-amniotic injection of HMGB1 induced the overexpression of several inflammatory genes in the fetal membranes, including a significant increase in *Ccl17* (Figure 4F&G). Conversely, M2 macrophage treatment exhibited a broad anti-inflammatory effect by downregulating the expression of *Tlr4*, *Cxcl1, Tnf*, and *Il1b* in the fetal membranes (Figure 4H&I). These findings indicate that M2 macrophages disrupt the inflammatory process of preterm labor in the fetal membranes.

**Figure 4.**
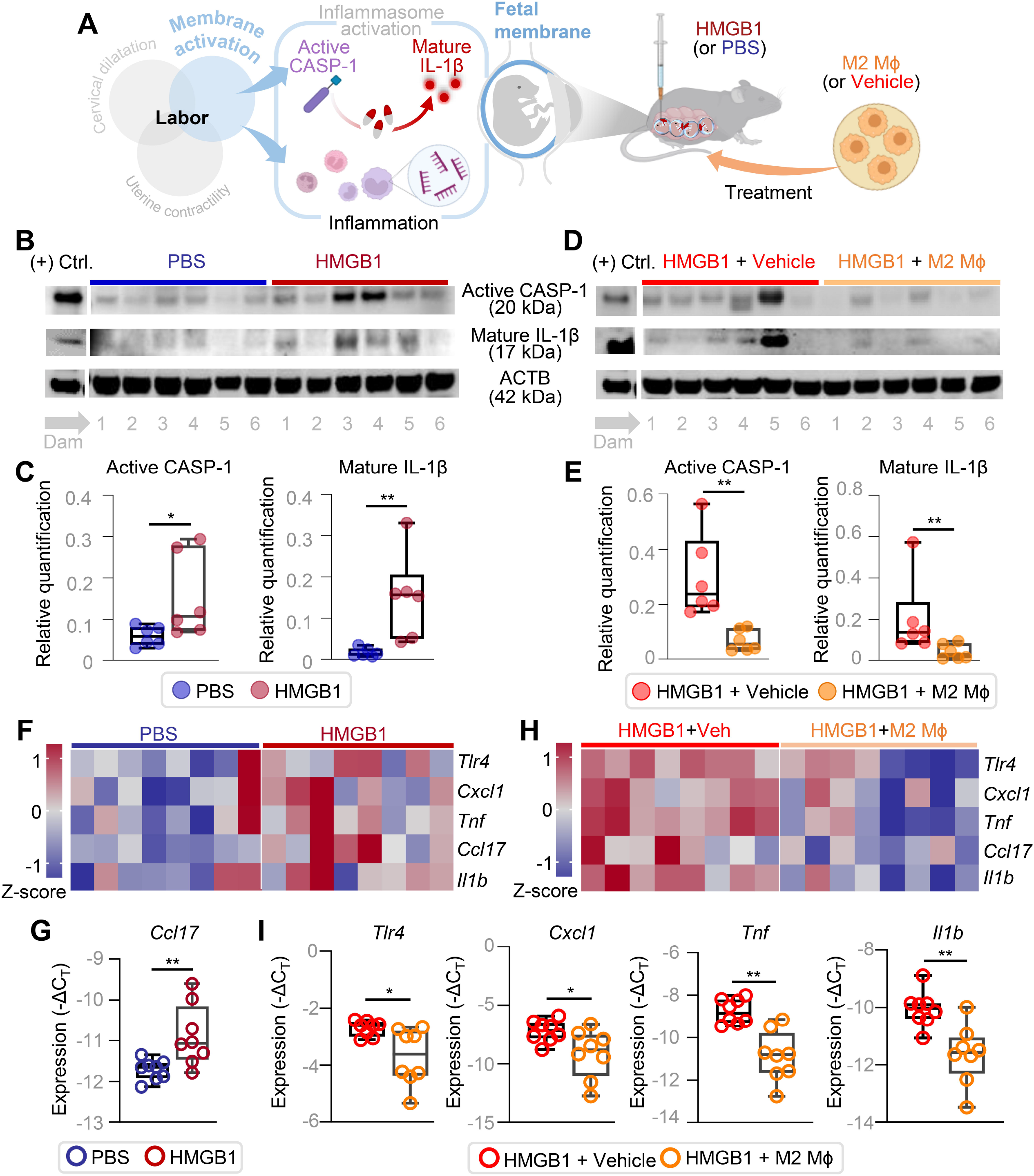
M2 macrophages inhibit inflammasome activation and regulate gene expression in the fetal membranes. **(A)** Dams received intra-amniotic injection of PBS (control) or HMGB1 on 14.5 days *post coitum* (dpc). M2 macrophages (M2 MΦ) or vehicle (PBS) were intravenously administered to dams on 13.5 dpc and 14.5 dpc followed by intra-amniotic injection of HMGB1 on 14.5 dpc. Tissue collection was performed at 72 h post-injection to collect the fetal membranes for immunoblotting and to determine gene expression. **(B)** Immunoblotting of active caspase (CASP)-1, mature IL-1β, and β-actin (ACTB) in the fetal membranes of PBS- or HMGB1-injected dams. Representative immunoblot images depict 6 samples per group in each gel. **(C)** Protein quantification of active CASP-1 and mature IL-1β (both normalized by ACTB) in the fetal membranes of PBS- or HMGB1-injected dams (n = 6 per group). **(D)** Immunoblotting of active CASP-1, mature IL-1β, and ACTB in the fetal membranes from HMGB1+Vehicle or HMGB1+M2 MΦ dams. Representative immunoblot images depict 6 samples per group in each gel. **(E)** Relative quantification of active CASP-1 and mature IL-1β (both normalized by ACTB) in the fetal membranes of HMGB1+Vehicle or HMGB1+M2 MΦ dams (n = 6 per group). **(F)** Representative heatmaps displaying the expression of key inflammatory genes in the fetal membranes of PBS-(n = 8) or HMGB1-injected (n = 8) dams. **(G)** Expression of *Ccl17* in the fetal membranes of PBS- or HMGB1-injected dams. Data are shown as boxplots where midlines indicate medians, boxes denote interquartile ranges, and whiskers indicate the minimum/maximum range. P-values were determined using the two-tailed Mann-Whitney U-test. **(H)** Representative heatmaps displaying the expression of key inflammatory genes in the fetal membranes of HMGB1+Vehicle (n = 8) or HMGB1+M2 MΦ (n = 8) dams. **(I)** Expression of *Tlr4*, *Cxcl1, Tnf*, and *Il1b* in the fetal membranes of HMGB1+Vehicle or HMGB1+M2 MΦ dams. Data are shown as boxplots where midlines indicate medians, boxes denote interquartile ranges, and whiskers indicate the minimum/maximum range. P-values were determined using the two-tailed Mann-Whitney U-test. *p < 0.05; **p < 0.01.

In the uterine tissues, intra-amniotic injection of HMGB1 induced inflammasome activation (Supplementary Figure 3A-H), but this was not significantly reduced by treatment with M2 macrophages. However, HMGB1 failed to induce a significant inflammatory response in uterine tissues, and treatment with M2 macrophages had no significant effect (Supplementary Figure 4A&B). These findings suggest that HMGB1-induced preterm labor in mice did not trigger a strong inflammatory response in uterine tissues, rendering M2 macrophage treatment unnecessary for preventing preterm birth.

In addition to the fetal membranes and uterus, the complex pathway of parturition also involves the initiation of an immune response in the decidual tissues^3,49,53^. However, we have demonstrated that such an inflammatory response is not associated with inflammasome activation^32,33,54^. Through inflammatory gene profiling, we found that HMGB1 did not strongly induce an inflammatory response in the decidua; yet, M2 macrophage treatment downregulated several inflammatory genes (Supplementary Figure 5A&B). This finding show that, even in the absence of abnormal decidual inflammation, M2 macrophages restrict the labor-associated inflammatory response at the maternal-fetal interface.

Together, these findings demonstrate that M2 macrophage treatment prevents preterm birth induced by *in utero* sterile inflammation by interfering with the inflammatory processes required to activate the pathway of preterm labor in the fetal membranes and maternal-fetal interface.

### M2 macrophage treatment mitigates inflammation in fetal and neonatal tissues

Given the robust homeostatic effects of M2 macrophages in the amniotic cavity and fetal membranes, we next investigated their ability to penetrate other fetal organs. We utilized a model wherein M2 macrophages derived from donor CD45.1^+^ mice were transferred to recipient CD45.2^+^ dams injected with HMGB1. Afterwards, maternal blood, maternal and fetal tissues, and amniotic fluid were collected to track the migration of transferred cells (Supplementary Figure 6A). Transferred M2 macrophages retained their M2 phenotype (data not shown), as previously reported^43^. Representative flow cytometry dot plots display the proportions of CD45.1^+^ M2 macrophages detected in maternal and fetal compartments (Supplementary Figure 6B). Quantification of transferred CD45.1^+^ M2 macrophages revealed distinct kinetic patterns across the maternal and fetal compartments (Supplementary Figure 6C&D). CD45.1^+^ M2 macrophages were highest in the maternal blood and lung at 2 hours post-injection, decreasing to negligible levels by 12 hours. Conversely, a gradual accumulation of CD45.1^+^ M2 macrophages was observed in the uterus, while modest numbers were consistently detected in the decidua at each time point (Supplementary Figure 6C). In the fetal compartments, CD45.1^+^ M2 macrophages were abundant in the placenta at 2 hours and declined over time, similar to the pattern in maternal blood, suggesting localization in the intervillous space rather than the parenchyma (Supplementary Figure 6D). A small number of CD45.1^+^ M2 macrophages accumulated in the fetal membranes over time, with a few reaching the amniotic cavity. In contrast, these cells were scarcely detected in the fetal intestine and lung at 2 hours, becoming negligible by 6 and 12 hours. The transient presence of M2 macrophages in the fetal lung and intestine is likely due to exposure to amniotic fluid, as evidenced by their absence in the unexposed fetal liver at any time point (Supplementary Figure 6D). These findings demonstrate that adoptively transferred M2 macrophages primarily migrate to intrauterine compartments, with limited penetration into fetal compartments.

Therefore, it is likely that M2 macrophages exert their homeostatic effects on the fetus through indirect mechanisms by downregulating the inflammatory response in the amniotic cavity and surrounding fetal membranes (Figures 3 and 4) and reducing inflammation in the placenta. We then evaluated the effect of M2 macrophage treatment on the placenta. The intra-amniotic injection of HMGB1 triggered the upregulation of *Tlr4*, *Gja1*, and *Ptgs2*, along with a non-significant increase in the expression of other genes, including *Ccl17* and *Nfkb2*, in the placenta (Supplementary Figure 7A). Treatment with M2 macrophages downregulated the expression of *Ccl17* and *Nfkb2* (Supplementary Figure 7B). Hence, M2 macrophages effectively dampen placental inflammation as part of their beneficial effects *in utero*.

Next, we focused on the fetal lung since preterm neonates born to women with intra-amniotic inflammation are at high risk of developing bronchopulmonary dysplasia^13–15^. In the fetal lung (Figure 5A), intra-amniotic injection of HMGB1 resulted in the upregulation of *Il6*, *Tnfrsf1a*, *Il33*, *Nlrp6*, and *Tlr9* (Figure 5B&C). M2 macrophage treatment downregulated the expression of *Tnfrsf1a* and *Nod1* (Figure 5D&E). Next, we explored whether the homeostatic effects of M2 macrophages were evident in the neonatal lung by evaluating inflammatory gene expression in this tissue. Neonates born to dams treated with M2 macrophages exhibited a diminished inflammatory profile in the neonatal lung, characterized by reduced expression of *Il6*, *Ccl2*, *Socs3*, and *Cxcl1* in comparison to offspring from untreated dams (Figure 5F&G). These findings indicate that M2 macrophages dampen inflammatory responses in the fetal lung, with sustained effects in the neonatal lung. This underscores the protective role of these cells in mitigating *in utero* sterile inflammation.

**Figure 5.**
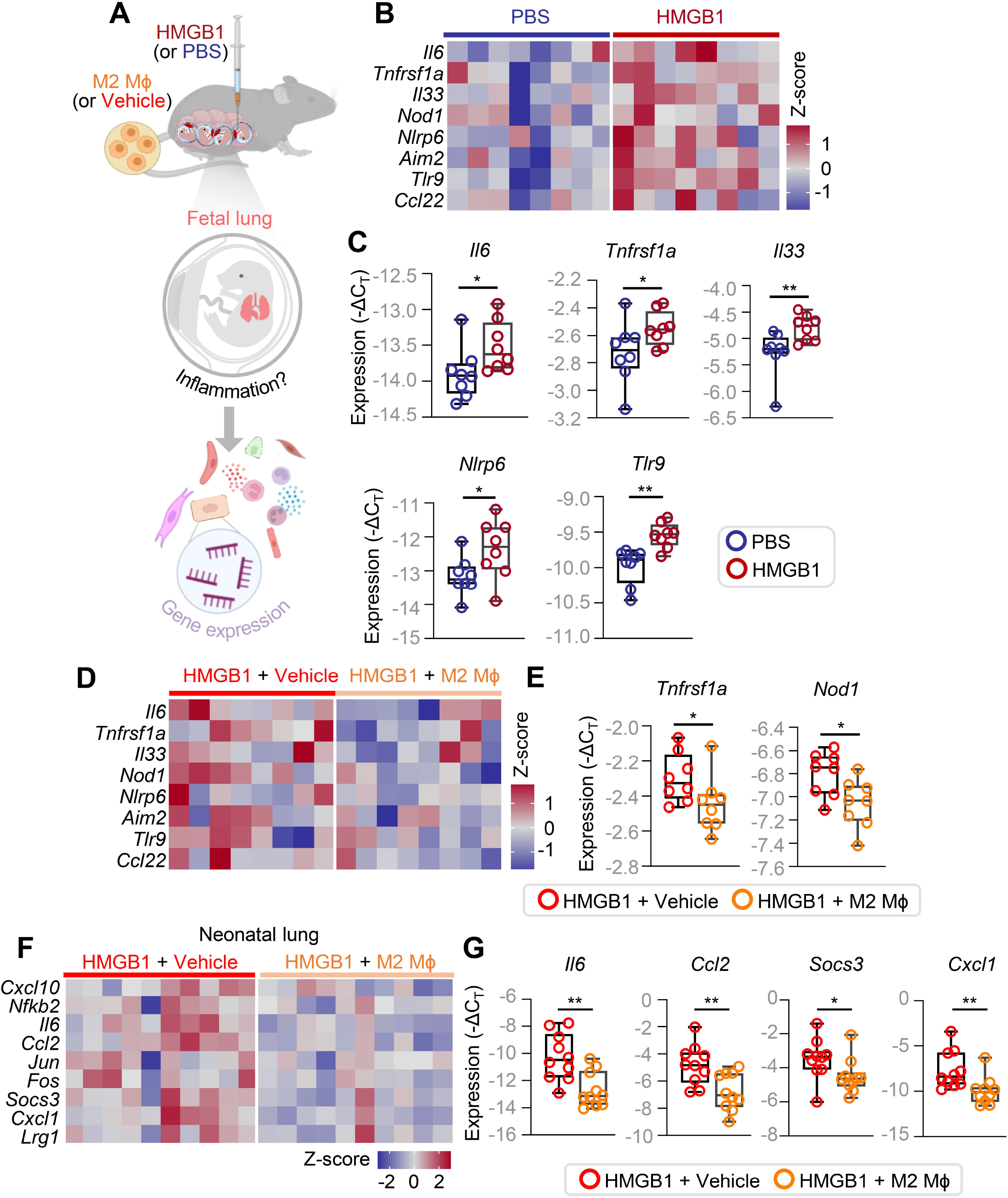
M2 macrophages ameliorate fetal and neonatal lung inflammation induced by *in utero* sterile inflammation. **(A)** Dams received intra-amniotic injection of PBS (control) or HMGB1 on 14.5 days *post coitum* (dpc). M2 macrophages (M2 MΦ) or vehicle (PBS) were intravenously administered to dams on 13.5 dpc and 14.5 dpc followed by intra-amniotic injection of HMGB1 on 14.5 dpc. Tissue collection was performed at 72 h post-injection to collect the fetal lung for gene expression. **(B)** Representative heatmaps displaying the expression of inflammatory genes in the fetal lung of PBS-(n = 8) or HMGB1-injected (n = 8) dams. **(C)** Expression of *Il6*, *Tnfrsf1a*, *Il33*, *Nlrp6*, and *Tlr9* in the fetal lung of PBS- or HMGB1-injected dams. Data are shown as boxplots where midlines indicate medians, boxes denote interquartile ranges, and whiskers indicate the minimum/maximum range. P-values were determined using the two-tailed Mann-Whitney U-test. **(D)** Representative heatmaps displaying the expression of inflammatory genes in the fetal lung of HMGB1+Vehicle (n = 8) or HMGB1+M2 MΦ (n = 8) dams. **(E)** Expression of *Tnfrsf1a* and *Nod1* in the fetal lung of HMGB1+Vehicle or HMGB1+M2 MΦ dams. **(F)** The lung was collected at 14-16 days of life from neonates born to HMGB1+Vehicle or HMGB1+M2 MΦ dams to evaluate gene expression as shown in the representative heatmap. **(G)** Expression of *Il6*, *Ccl2*, *Socs3*, and *Cxcl1* in the neonatal lung (n = 10 per group). Data are shown as boxplots where midlines indicate medians, boxes denote interquartile ranges, and whiskers indicate the minimum/maximum range. P-values were determined using the two-tailed Mann-Whitney U-test. *p < 0.05; **p < 0.01.

In addition to the lungs, the fetal intestine is also exposed to amniotic fluid, putting neonates born to women with intra-amniotic inflammation at high risk of developing necrotizing enterocolitis^16,17^. Therefore, we next explored inflammatory gene expression in the fetal intestine following an intra-amniotic injection of HMGB1. This revealed alterations in inflammatory gene expression (Supplementary Figure 8), consistent with our previous reports on *in utero* inflammation^55,56^. Unlike in the fetal lung, the inflammatory milieu in the fetal intestine was downregulated, suggesting that *in utero* sterile inflammation exerts different effects across fetal organs. Since we previously showed that treatment with an anti-inflammatory drug did not exert a significant impact on the fetal intestine exposed to *in utero* sterile inflammaiton^55^, we shifted our focus to investigate the effects of M2 macrophage treatment on the neonatal intestine (Figure 6A). Treatment with M2 macrophages boosted the neonatal gut inflammatory response dampened by *in utero* exposure to HMGB1 (Figure 6B, D, F). Specifically, M2 macrophage treatment upregulated the expression of *Aim2, Cd68, Tnf, Rgs4, Ccl5,* and *Ccl17* in the small intestine; *Casp1* and *Ccl17* in the cecum; and *Il1a, Cd68, Casp1, Jun, Kdm6b,* and *Ccl17* in the colon (Figure 6C, E, G) of neonates. These data show that treatment with M2 macrophages enhances the neonatal gut inflammatory profile, which is otherwise suppressed by *in utero* exposure to sterile inflammation.

**Figure 6.**
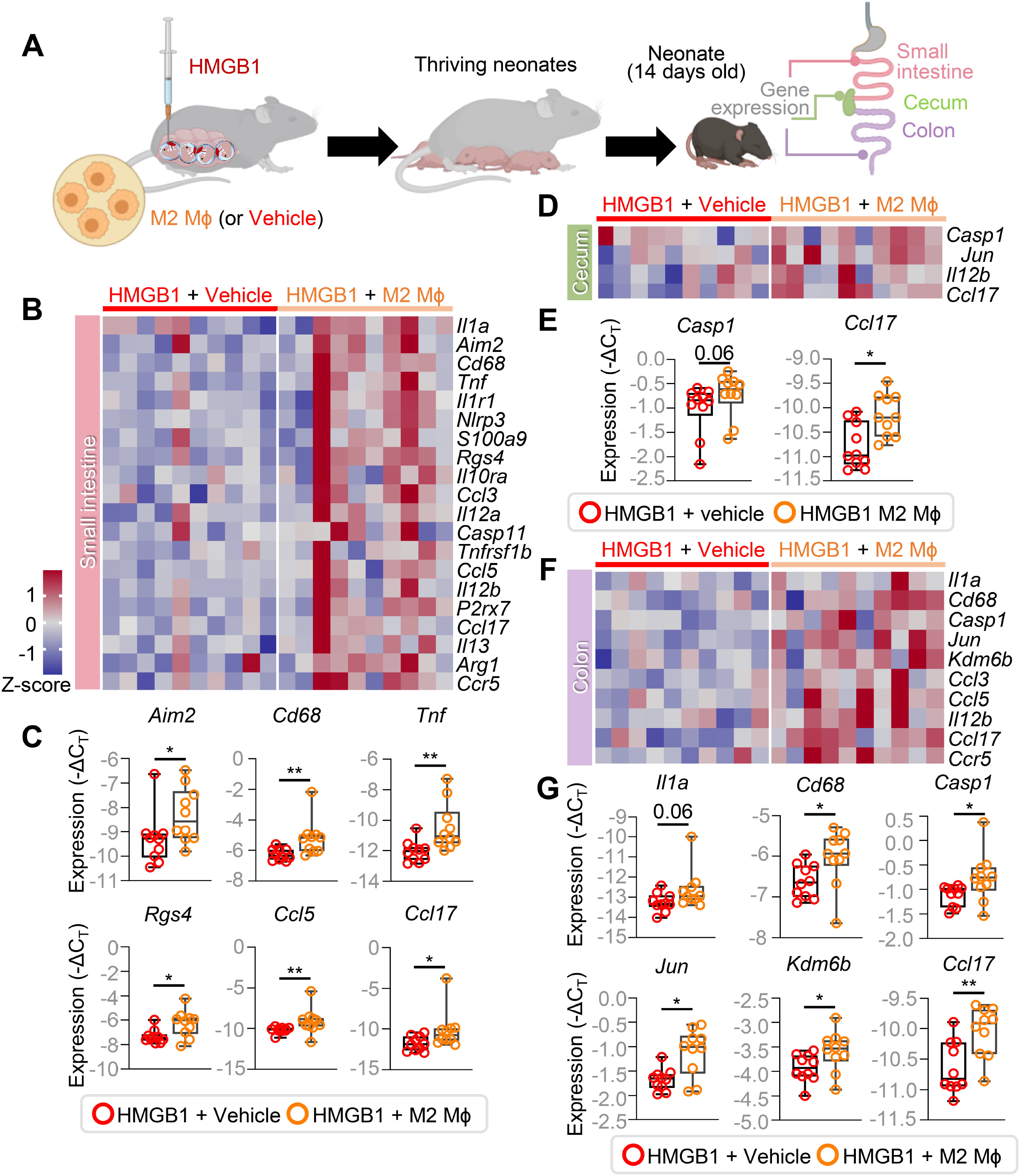
M2 macrophages regulate neonatal gut inflammation resulting from *in utero* sterile inflammation. **(A)** M2 macrophages (M2 MΦ) or PBS (Vehicle) were intravenously administered to dams on 13.5 dpc and 14.5 dpc followed by intra-amniotic injection of HMGB1 on 14.5 dpc. After delivery, neonates were monitored until 14-16 days of age, after which the small intestine, cecum, and colon were collected to determine gene expression. Representative heatmaps display gene expression in the **(B)** neonatal small intestine, **(D)** cecum, and **(F)** colon of neonates born to HMGB1+Vehicle (n = 10 neonates) or HMGB1+M2 MΦ (n = 10 neonates) dams. The expression of specific genes in the **(C)** neonatal small intestine (*Aim2, Cd68, Tnf, Rgs4, Ccl5, Ccl17*), **(E)** cecum (*Casp1* and *Ccl17*), and **(G)** colon (*Il1a, Cd68, Casp1, Jun, Kdm6b, Ccl17*) are shown as box plots. P-values were determined using the two-tailed Mann-Whitney U-test. Data are shown as boxplots where midlines indicate medians, boxes denote interquartile ranges, and whiskers indicate the minimum/maximum range. *p < 0.05; **p < 0.01. See also Figures S8-S10.

We recently demonstrated that *in utero* sterile inflammation leads to neonatal gut dysbiosis^55^. Therefore, we proceeded to evaluate whether treatment with M2 macrophages could mitigate this dysbiosis by performing 16S rRNA gene sequencing of the small intestine, cecum, and colon from neonates born to dams intra-amniotically injected with HMGB1 with or without M2 macrophage treatment (Supplementary Figure 9A). Alpha diversity metrics of the microbiome indicated no differences in the community evenness of the small intestine, cecum, or colon between groups (data not shown). When examining beta diversity, principal coordinate analysis (PCoA) revealed that the neonatal microbiomes were strongly clustered by treatment group (Supplementary Figure 9B) and underwent consistent changes in structure and Firmicutes/Bacteroidetes ratio (Supplementary Figure 9C-E). In the small intestine, M2 macrophage treatment was associated with a shift in the microbiome structure (Supplementary Figures 9D and 10A&B) characterized by decreased abundance of *Rodentibacter* (ASV-5) and *Ruminiclostridium* (ASV-17) (Supplementary Figure 10C). Similar patterns were also observed in the cecum (Supplementary Figures 9D and 10D&E), in which a decreased abundance of *Ruminiclostridium* (ASV-17), Lachnospiraceae (ASV-24), and *Lachnoclostrium* (ASV-15) was observed alongside an increased abundance of Lachnospiraceae (ASV-39) (Supplementary Figure 10F). Consistently, in the colon, the microbiome structure was modified by M2 macrophage treatment (Supplementary Figures 9D and 10G&H), resulting in decreased abundance of *Ruminiclostridium* (ASV-17), Lachnospiraceae (ASV-24), and *Lachnoclostrium* (ASV-15) (Supplementary Figure 10I). Treatment with M2 macrophages alone did not alter the alpha or beta diversity of the neonatal gut microbiome (Supplementary Figure 9F-H). Thus, prenatal treatment with M2 macrophages can change neonatal gut dysbiosis induced by *in utero* exposure to sterile inflammation.

Collectively, the data demonstrate that prenatal treatment with M2 macrophages dampens fetal tissue inflammation, thereby preventing subsequent neonatal organ inflammatory injury and gut microbiome dysbiosis. These findings suggest potential mechanisms through which this cellular therapy improves neonatal outcomes following exposure to *in utero* sterile inflammation.

### M2 macrophage treatment improves neonatal ability to fight Group B *Streptococcus* infection following *in utero* sterile inflammation

Thus far, we have demonstrated that M2 macrophage treatment enhances survival and alleviates inflammation in neonates exposed to *in utero* sterile inflammation. Therefore, we last assessed whether these changes would result in improved neonatal ability to combat systemic infection with Group B *Streptococcus* (GBS), a microbe commonly associated with neonatal infection and sepsis^57^. At two weeks of age, control neonates (i.e., pups born to dams without any treatment) and neonates exposed *in utero* to HMGB1 alone, HMGB1 with vehicle treatment, or HMGB1 with M2 macrophage treatment received intraperitoneal injection of GBS (Figure 7A). *In utero* exposure to HMGB1 caused low neonatal survival rates within the first five days post-GBS infection (Figure 7B). Notably, prenatal treatment with M2 macrophages bolstered neonatal survival to be comparable to controls (Figure 7B), indicating improved capacity to fight GBS infection. *In utero* exposure to HMGB1 also negatively impacted neonatal weight, which was significantly reduced compared to controls at five days post-infection (Figure 8cC). Prenatal treatment with M2 macrophages partially rescued neonatal weight but did not fully restore it to the control trajectory (Figure 7C), a phenomenon that requires further investigation. The improved neonatal survival and growth driven by M2 macrophage treatment were accompanied by reduced incidence of hypothermia (considered the murine equivalent of fever^58,59^) upon GBS infection (Supplementary Figure 11). Thus, prenatal treatment with M2 macrophages bolsters the capacity to clear pathogenic bacteria in neonates compromised by exposure to *in utero* sterile inflammation.

**Figure 7.**
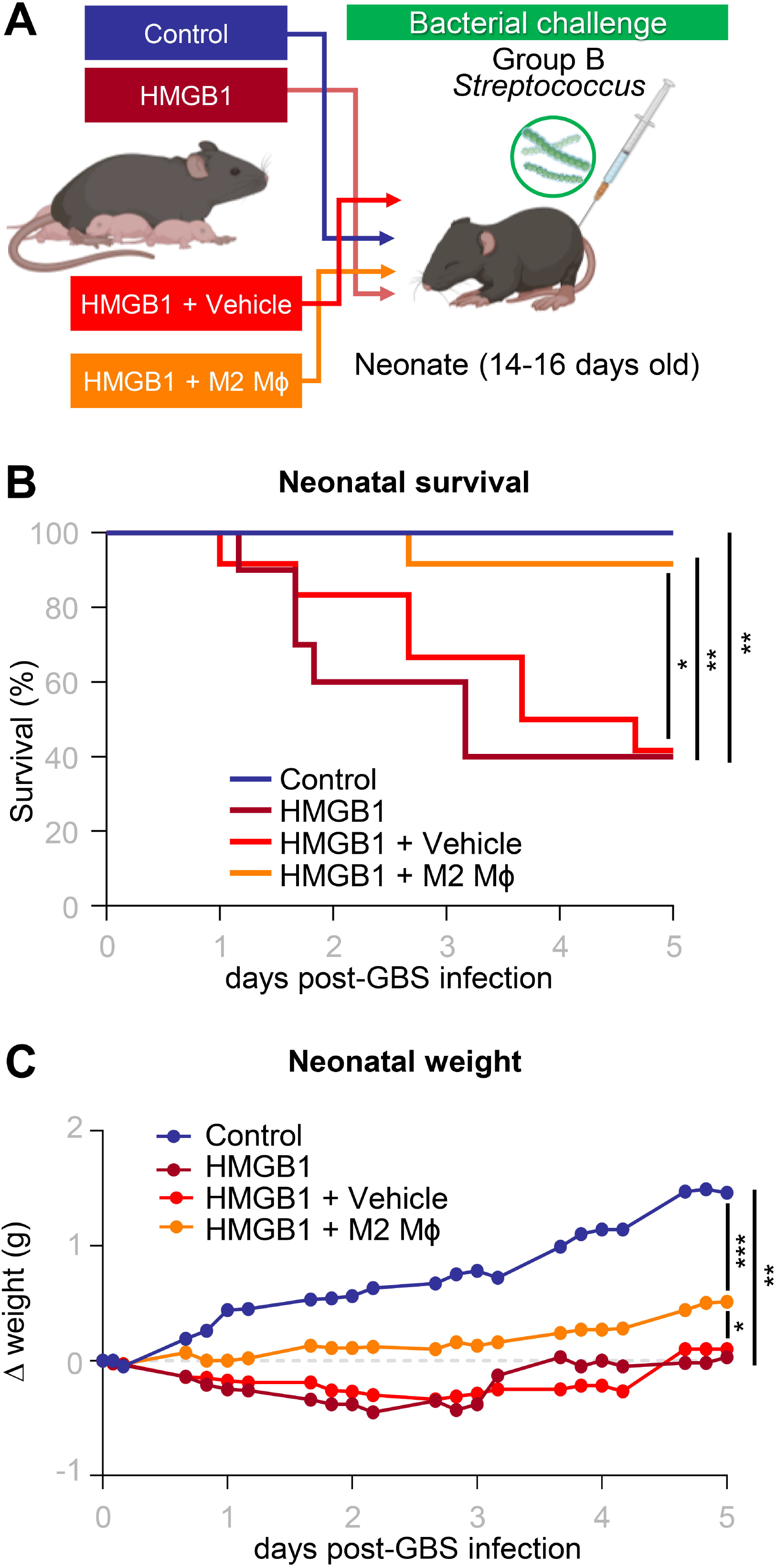
M2 macrophages improve neonatal immunocompetence. **(A)** M2 macrophages (M2 MΦ) or PBS (Vehicle) were intravenously administered to dams on 13.5 dpc and 14.5 dpc followed by intra-amniotic injection of HMGB1 on 14.5 dpc. Pregnant dams without any treatment or injected with HMGB1 alone were also included. After delivery, surviving neonates were monitored until 14-16 days of age, after which they received intraperitoneal injection of Group B *Streptococcus* (GBS) and were observed for five days. **(B)** The survival rates of GBS-infected neonates born to untreated (n = 10 neonates), HMGB1-injected (n = 10 neonates), HMGB1+Vehicle (n = 12 neonates), or HMGB1+M2 MΦ (n = 12 neonates) dams over the five days post-challenge are displayed as Kaplan-Meier survival curves. P-values were determined using the Gehan-Breslow-Wilcoxon test. **(C)** Mean weights of GBS-infected neonates over the five days post-challenge. P-values for comparing the mean weight at the end of day five (final data point in plot) were determined using the two-tailed Mann-Whitney U-test. *p < 0.05; **p < 0.01; ***p < 0.001. See also Figure S11.

## DISCUSSION

The lack of knowledge regarding specific molecular targets in women with *in utero* sterile inflammation has made the investigation of useful treatment options challenging. In the current study, we provide mechanistic evidence for M2 macrophage treatment as a cellular therapy that can prevent preterm birth resulting from *in utero* sterile inflammation. The reasoning behind choosing M2 macrophages as an approach to prevent preterm birth and the mechanisms through which they exert these effects are explained below. *First*, our prior studies have indicated that both physiological and pathological labor are accompanied by a shift in macrophage polarization, primarily at the maternal-fetal interface, with a subset of these cells acquiring a pro-inflammatory M1-like phenotype^42,43^. Here, we build on this concept by showing that M2-like homeostatic macrophages of maternal origin are reduced in the myometrium and basal plate of women with term and preterm labor, respectively. This finding complements our previous targeted approaches, which showed a reduction of decidual M2 macrophages in women experiencing preterm labor and birth^43^. Hence, it is possible that homeostatic macrophages in the uterine and decidua tissues help maintain a favorable environment at the maternal-fetal interface and surrounding tissues, thereby dampening inflammatory signaling associated with the premature onset of labor. The latter concept is supported by our prior observations that the depletion of maternal macrophages can result in preterm labor and birth^43^. *Second*, labor involves coordinated activation across multiple maternal and fetal tissues, conventionally referred to as the common pathway of parturition^47–50^. Herein, we demonstrate that M2 macrophages infiltrate the maternal-fetal interface, interfere with the common pathway of labor in the fetal membranes and decidua, and dampen fetal inflammation mediated by the placenta and amniotic cavity. In line with this, M2 macrophages express immunomodulatory factors, such as arginase-1 and IL-10, that can modify the tissue microenvironment and prevent local immune activation^60,61^. Indeed, in a previous study using an animal model of LPS-induced intra-amniotic inflammation, we found that treatment with M2 macrophages increased concentrations of IL-10 in both maternal circulation and amniotic fluid, along with a reduction in inflammatory mediators at the maternal-fetal interface^43^, providing further evidence that homeostatic macrophages target both maternal and fetal components in the process of preterm labor. *Third*, we previously demonstrated that the inflammasome is a key pathway involved in premature labor induced by microbes^54,62^ or alarmins^32–34,63^. As a proof of concept, we administered a specific inhibitor of the NLRP3 inflammasome (i.e., MCC950)^64^ to dams injected with the alarmin S100B and observed reduced rates of preterm birth and adverse neonatal outcomes^32^. While NLRP3 inhibitors hold promise, further research is required to comprehensively assess their effects on offspring before considering their use during pregnancy. The current study demonstrates that M2 macrophages suppress inflammasome activation in the amniotic cavity and fetal membranes, offering an alternative strategy for modulating this signaling pathway.

*In utero* sterile inflammation is not only detrimental due to its short-term outcomes, such as preterm birth, but is importantly linked to long-term negative consequences for the offspring as indicated by human data^7,11^ and animal studies^27–29,32,34,55,63,65^. Such consequences are not entirely the result of prematurity, as neonates born at term to HMGB1-injected dams still had reduced survival compared to those that received treatment with M2 macrophages. Thus, even a relatively brief exposure to *in utero* sterile inflammation imprints fetal tissue alterations that can be carried over to neonatal life. This concept is supported by previous research demonstrating that prenatal treatments or exposures during gestation can induce permanent changes in neonatal life^55,66–68^. Of interest, recent research employing a transient gestational infection model revealed the role of the IL-6/Th17 axis in augmenting offspring gut immunity, indicating that prenatal immunological events can induce long-lasting effects^67^. Transient exposure to a maternally-derived inflammatory cytokine, IL-6, was similarly explored in pregnant mice and found to result in permanent alteration of neuronal gene programs, providing mechanistic evidence for how prenatal inflammation drives long-term changes in neonatal neurodevelopment^68^. Strikingly, in the current study, we report that prenatal treatment with M2 macrophages mitigates the negative consequences of *in utero* sterile inflammation, enhancing survival, reversing growth restriction, and overall improving neonatal infection resistance. Therefore, the current study supports two key concepts: firstly, acute insults *in utero* can disrupt developmental programming, resulting in impaired immunocompetence in the offspring, the full extent and significance of which remain poorly understood; and second, the prenatal cellular therapy using M2 macrophages demonstrated herein exerts substantial homeostatic effects, effectively countering such alterations.

Proper formation of the gut microbiota during early life is essential for the development of a healthy adult gut^69^. Indeed, multiple inflammatory, metabolic, neurologic, cardiovascular, and gastrointestinal diseases have been linked to disruption of the microbiota during early life^70^. Importantly, the risk of gut dysbiosis is increased in preterm neonates compared to those delivered at term^69^, and thus the prevention or treatment of premature labor leading to preterm birth, particularly in the context of *in utero* sterile inflammation, is essential to ensure the development of a healthy gut microbiota and avoid long-term consequences. Several prenatal events have been studied and found to contribute to microbiota dysbiosis in human neonates, including maternal diet and antibiotic use, among others^69,71^. Specifically, the combined presence of acute histologic chorioamnionitis/funisitis was linked to altered microbial composition in neonatal fecal samples, including modified abundance of *Bacteroidetes* and *Fusobacteria*^72^. Importantly, neonates affected by acute histologic chorioamnionitis alone or acute histologic chorioamnionitis/funisitis had higher incidence of late-onset sepsis/death, thereby providing an association between microbiome alterations and short-term adverse events^72^. Consistent with these studies, we demonstrated herein that adverse neonatal outcomes resulting from *in utero* sterile inflammation are similarly linked to alterations in gut microbiome composition, and that prenatal treatment with M2 macrophages improves these aberrations. Although the current study did not continuously examine the neonatal gut microbiome from birth to the weaning period, our results indicate that treatment with M2 macrophages promotes the proper maturation of a "healthy" gut microbiota, which can contribute to reducing disease risk in neonatal life.

It is worth acknowledging that there are inherent differences between humans and murine models. Key physiological factors such as gestational length, placental structure, immune system regulation, and hormonal regulation vary between humans and mice, as has been extensively described^73–75^. Therefore, we consider that any interventions that prove effective in murine models, such as the M2 macrophages demonstrated in the current study, require additional validation in models that more closely resemble human, such as non-human primates, to more firmly establish their clinical relevance during pregnancy.

Collectively, the data presented herein link a reduction in maternal homeostatic macrophages to human labor and mechanistic evidence in mice demonstrating that M2 macrophages can treat *in utero* sterile inflammation, thereby preventing preterm birth and adverse neonatal outcomes. Specifically, treatment with M2 macrophages mitigates inflammatory responses in the amniotic cavity and surrounding fetal membranes by inhibiting inflammasome activation and halting inflammatory processes in the common pathway of labor, thereby extending gestational length and preventing preterm birth. Importantly, M2 macrophage treatment not only diminishes inflammation in the placenta as well as the fetal and neonatal lung but also enhances neonatal intestinal responses. Neonates born to dams prenatally treated with M2 macrophages demonstrate a distinct gut microbiome compared to those born from dams with *in utero* sterile inflammation, alongside an improved ability to clear bacterial infections. In conclusion, these findings provide mechanistic evidence that M2 macrophages can serve as a cellular approach not only to prevent premature labor leading to preterm birth but also to enhance fetal and neonatal immunity following *in utero* exposure to sterile inflammation.

## Supporting information

Key Resource Table

## Acknowledgements

This research was supported by the Perinatology Research Branch, Division of Obstetrics and Maternal-Fetal Medicine, Division of Intramural Research, *Eunice Kennedy Shriver* National Institute of Child Health and Human Development, National Institutes of Health, U.S. Department of Health and Human Services (NICHD/NIH/DHHS) under Contract No. HHSN275201300006C. This research was also supported by the Wayne State University Perinatal Initiative in Maternal, Perinatal and Child Health. R.R. has contributed to this work as part of his official duties as an employee of the United States Federal Government. The funders had no role in study design, data collection and interpretation, or the decision to submit the work for publication. Figures include art created with BioRender.com

## Author contributions

Conceptualization: N.G.-L.

Methodology: V.G.-F., Z.L., Y.X., N.G.-L.

Validation: V.G.-F., Z.L., Y.X., N.G.-L.

Formal analysis: V.G.-F., Z.L., R.R., R.P.-R., Y.X., D.L., J.G., A.D.W., M.F.-J., J.J.P., N.G.-L.

Investigation: V.G.-F., Z.L., R.P.-R., Y.X., D.M., D.L., J.G., A.D.W., M.F.-J., J.J.P., N.G.-L.

Resources: R.R., K.R.T., N.G.-L.

Data curation: V.G.-F., Z.L., R.P.-R., Y.X., A.D.W.

Writing – original draft: V.G.-F., D.M., J.G., N.G.-L.

Writing – review & editing: V.G.-F., Z.L., R.R., R.P.-R., Y.X., D.M., D.L., J.G., A.D.W., M.F.-J., J.J.P., K.R.T., N.G.-L.

Visualization: V.G.-F., Z.L., J.G., A.D.W., N.G.-L.

Supervision: R.R., R.P.-R., K.R.T., N.G.-L.

Project administration: N.G.-L.

Funding acquisition: R.R., N.G.-L.

## STAR METHODS

### KEY RESOURCES TABLE

(Submitted separately)

## RESOURCE AVAILABILITY

### Lead Contact

Further information and requests for resources and reagents should be directed to and will be fulfilled by the lead contact, Nardhy Gomez-Lopez.

### Materials Availability

This study did not generate new unique reagents.

### Data and Code Availability

Single-cell RNA-sequencing data were previously reported^44–46^ and are deposited in the NIH dbGAP database (phs001886.v5.p1). 16S rRNA gene sequencing files and associated metadata have been uploaded to the National Center for Biotechnology Information’s Sequence Read Archive (PRJNA925285). All other data are presented within the current manuscript and/or its supplementary materials.

## EXPERIMENTAL MODEL AND SUBJECT DETAILS

C57BL/6 (strain # 000664) and B6 CD45.1 (B6.SJL-*Ptprc^a^ Pepc^b^*/BoyJ; strain # 002014) mice were purchased from The Jackson Laboratory (Bar Harbor, ME, USA). Mice were housed in the animal care facility at the C.S. Mott Center for Human Growth and Development at Wayne State University (Detroit, MI, USA) under a circadian cycle (light:dark = 12:12 h). Eight-to twelve-week-old females were mated with males of proven fertility. Females were checked daily between 8 and 9 a.m. to investigate the appearance of a vaginal plug, which indicated 0.5 days *post coitum* (dpc). Plugged females were then housed separately from the male mice and their weights were monitored daily. A weight gain of ≥ 2 grams by 12.5 dpc confirmed pregnancy. Mice were randomly assigned to study groups prior to the experiments described herein. Numbers of biological replicates are indicated in each figure caption. All procedures were approved by the Institutional Animal Care and Use Committee (IACUC) at Wayne State University under Protocol Nos. A-07-03-15, 18-03-0584, and 21-04-3506.

## METHOD DETAILS

### Analysis of scRNA-seq data

#### Data normalization and pre-processing

Data from three of our previously generated human scRNA-seq datasets^44–46^ were combined and re-analyzed to specifically investigate macrophage populations at the maternal-fetal interface. All datasets were pre-processed and normalized with a comparable pipeline. In brief, sequencing data were processed using Cell Ranger version 4.0.0 (10x Genomics). The fastq files were aligned using kallisto^76^, and bustools^77^ was used to summarize the cell/gene transcript counts in a matrix for each sample. In parallel, “cellranger counts” was also used to align the scRNA-seq reads using the STAR aligner^78^ to produce the bam files necessary for demultiplexing the individual of origin based on genotype information using souporcell^79^ and demuxlet^80^. Quality metrics were calculated and each library was determined to be of excellent quality based on 10X Genomics recommendations. Any droplet/GEM barcode assigned to a double or ambiguous cell in demuxlet or souporcell was filtered, and only cells that could be assigned to a pregnancy case were kept. Furthermore, any cells with less than 100 or more than 10,000 genes were filtered out, as well as those with > 25% mitochondrial reads. Cells annotated as macrophages (details regarding the annotation process can be found in the original publications^44–46^) were extracted and combined into a new Seurat object.

All count data matrices were normalized and combined using the “NormalizedData” “FindVariableFeatures” and “ScaleData” methods implemented in the Seurat package in R^81,82^. Next, the Seurat “RunPCA” function was applied to obtain the first 50 principal components, and the different libraries were integrated and harmonized using the Harmony package in R^83^ while accounting for library of origin as a potential batch effect. The top 30 harmony components were then processed using the Seurat “runUMAP” function to embed and visualize the cells in a two-dimensional map via the Uniform Manifold Approximation and Projection for Dimension Reduction (UMAP) algorithm. The “FindClusters” function with a resolution of 0.1 was used to cluster the single cells into seven distinct clusters. The two most abundant macrophage cell type clusters (M1 and M2) were found to be equivalent to the previously annotated clusters Macrophage-1 and Macrophage-2, respectively that we have previously reported^44–46^. The proportions of specific macrophage clusters within a tissue were compared between groups using Mann-Whitney <u>U</u>-tests.

#### Gene ontology

The differential expression of selected marker genes for each cell type/cluster were identified using the Wilcoxon Rank Sum test implemented by FindAllMarkers function from Seurat. For this analysis, we compared each cluster to all cell types. The clusterProfiler^84^ was used to perform Over-Representation Analysis (ORA) based on the Gene Ontology (GO) using “enrichGO” function. We first compared all clusters to each other using “compareCluster” and reported the top terms for each cluster. Since many of these terms were shared, we then focused on the M2 cluster and compared the list of unique genes identified with a FDR of 5% for this population against the universe of all genes expressed in macrophages. Only ORA results that were significant after correction were reported with *q* < 0.05 being considered statistically significant.

### Intra-amniotic administration of HMGB1

Ultrasound-guided intra-amniotic injection of HMGB1 was performed as previously reported^27–29^. Briefly, dams were anesthetized on 14.5 dpc by inhalation of 2% isoflurane (Fluriso^TM^/Isoflurane, USP; VetOne, Boise, ID, USA) and 1–2 L/min of oxygen in an induction chamber, and a mixture of 1.5–2% isoflurane and 1.5–2 L/min of oxygen was used to maintain anesthesia. Mice were positioned on a heating pad and stabilized with adhesive tape, and fur was removed from the abdomen and thorax using Nair cream (Church & Dwight Co., Inc., Ewing, NJ, USA). Body temperature was detected with a rectal probe (VisualSonics, Toronto, ON, Canada) throughout the procedure, and respiratory and heart rates were monitored by electrodes embedded in the heating pad. An ultrasound probe was fixed and mobilized with a mechanical holder, and the transducer was slowly moved toward the abdomen^85^. The ultrasound-guided intra-amniotic injection of recombinant human HMGB1 (Biolegend, San Diego, CA, USA) at a μL of sterile 1X phosphate-buffered saline (PBS; Life Technologies, Grand Island, NY, USA) was performed in each amniotic sac using a 30G needle (BD PrecisionGlide Needle; Becton Dickinson, Franklin Lakes, NJ, USA). After ultrasound completion, mice were placed under a heat lamp for recovery defined as when the mouse resumed normal activities, such as walking and interacting with its environment, which typically occurred within 10 min after removal from anesthesia. After recovery, dams were monitored via video camera (Sony Corporation, Tokyo, Japan) to observe pregnancy and neonatal outcomes.

### Isolation, differentiation, and adoptive transfer of M2 macrophages

Bone marrow cells were isolated, differentiated, and adoptively transferred following a previously established protocol^43^. Briefly, bone marrow cells were collected from female mice (12-16 weeks old) and treated with red blood cell lysis buffer (Ammonium Chloride Solution; Cat# 07800, Stem Cell Technologies; Vancouver, CA). Then, cells were resuspended and cultured in Iscove’s Modified Dulbecco’s Media (IMDM) medium (Thermo-Fisher Scientific; Waltham, MA, USA) supplemented with 10% fetal bovine serum (FBS; Invitrogen, Carlsbad, CA, USA), 1X antibiotic-antimycotic (Cat# 15240062; Thermo-Fisher Scientific), and 10 ng/ml recombinant macrophage colony-stimulating factor (M-CSF; Cat# 576402; BioLegend, San Diego, CA, USA) for 7 days. On day 7, the culture medium was replaced with fresh IMDM medium supplemented with 10% FBS, 1X antibiotic-antimycotic, 10 ng/ml of recombinant IL-4 (Cat# 574302, BioLegend), and 10 ng/ml of recombinant IL-13 (Cat# 575902, BioLegend). After 14-18 h, M2 macrophages were collected by washing with ice-cold PBS (Fisher Scientific Bioreagents, Fair Lawn, NJ, USA or Life Technologies Limited, Pailey, UK). The purity and phenotype of macrophages after M2 polarization were routinely checked throughout the course of the study, as we have previously reported^43^. Prior to injection, the viability of M2 macrophages was consistently confirmed to be >90%. Approximately 2-5 x 10^6^ cells were μL PBS for intravenous injection into dams on 13.5 and 14.5 dpc, prior to the administration of HMGB1 or PBS on 14.5 dpc. This timing was chosen so that the transferred M2 macrophages would have sufficient time to reach key maternal and fetal tissues during the critical window of inflammation resulting in preterm birth, as previously reported^43^.

### Perinatal outcomes

Gestational length was calculated as the duration of time from the presence of the vaginal plug (0.5 dpc) to the detection of the first pup in the cage bedding. Preterm birth was defined as delivery occurring before 18.75 dpc based on the lowest gestational age at delivery observed in the control group. The rate of preterm birth was calculated as the proportion of females delivering preterm out of the total number of mice per group. The rate of stillbirth was defined as the proportion of delivered pups found dead among the total number of pups. The rate of neonatal mortality was defined as the proportion of pups found dead among the total number of pups. Neonatal mortality rates and weights were calculated and recorded at postnatal weeks 1, 2, and 3.

### Sampling from dams intra-amniotically injected with HMGB1

Dams were euthanized at 24, 48, 72, and 96 h after intra-amniotic HMGB1 injection for tissue collection. The amniotic fluid was collected from each amniotic sac and centrifuged at 1,300 x g for 5 min at 4°C. The resulting supernatants were stored at -20°C until analysis. The maternal whole blood was collected from each dam, mixed with heparin (Cat# 2106-15VL; Sigma-Aldrich, St. Louis, MO, USA), and centrifuged at 800 x g for 10 min at 4°C. Collection of the uterus, decidua, placenta, fetal membrane, fetal lung, and fetal intestine was performed. The placentas and fetuses were imaged and weighed during tissue dissection. The collected tissues were snap-frozen in liquid nitrogen for immunoblotting or preserved in RNA*later* Stabilization Solution (Cat# AM7021; Invitrogen by Thermo-Fisher Scientific, Carlsbad, CA, USA), according to the manufacturer’s instructions.

### Determination of cytokine concentrations in amniotic fluid

Concentrations of IL-6, IL-1β, and TNF were determined in murine amniotic fluid using the U-PLEX Custom Biomarker Group 1 (ms) assay (Cat# K15069M-2; Meso Scale Discovery, Rockville, MD, USA), according to the manufacturer’s instructions. Plates were read using the MESO QuickPlex SQ 120 (Meso Scale Discovery) and analyte concentrations were calculated using the Discovery Workbench software v4.0 (Meso Scale Discovery). The sensitivities of IL-6, IL-1β, and TNF were 4.8 pg/ml, 3.1 pg/ml, and 1.3 pg/ml, respectively.

### RNA isolation, cDNA synthesis, and qPCR analysis of tissues from pregnant mice

Total RNA was isolated from the uterus, decidua, placenta, fetal membranes, fetal lung, and fetal intestine using QIAshredders (Cat# 79656; Qiagen, Germantown, MD, USA), RNase-Free DNase Sets (Cat# 79254; Qiagen), and RNeasy Mini Kits (Cat# 74106; Qiagen), according to the manufacturer’s instructions. The NanoDrop 8000 spectrophotometer (Thermo Scientific, Wilmington, DE, USA) and the Bioanalyzer 2100 (Agilent Technologies, Waldbronn, Germany) were used to evaluate RNA concentrations, purity, and integrity. SuperScript IV VILO Master Mix (cat# 11756050; Invitrogen by Thermo Fisher Scientific Baltics UAB, Vilnius, Lithuania) was used to synthesize complementary (c)DNA. Gene expression profiling of the tissues was performed on the BioMark System for high-throughput qPCR (Fluidigm, San Francisco, CA, USA) with the TaqMan gene expression assays (Applied Biosystems, Life Technologies Corporation, Pleasanton, CA, USA) listed in Supplementary Table 2. Negative delta cycle threshold (-ΔC_T_) values were determined using multiple reference genes (*Actb*, *Gapdh*, *Gusb*, and *Hsp90ab1*) averaged within each sample for contractility-associated and inflammatory genes. The -ΔC_T_ values were normalized by calculating the Z-score of each gene with the study groups. Heatmaps were created representing the Z-score of each -ΔC_T_ using GraphPad Prism (GraphPad, San Diego, CA, USA).

### Tissue lysate preparation and protein quantification

Snap-frozen fetal membrane and uterus were mechanically homogenized in PBS containing a complete protease inhibitor cocktail (Cat# 11836170001; Roche Applied Sciences, Mannheim, Germany) for 10 min. The resulting lysates were centrifuged at 16,100 x g (maximum speed) for 5 min at 4°C. The total protein concentrations in amniotic fluid (supernatant) and tissue lysate samples were tested by the Pierce BCA Protein Assay Kit (Cat# 23225; Pierce Biotechnology, Thermo-Fisher Scientific, Inc., Rockford, IL, USA) prior to immunoblotting. Cell lysates or concentrated culture medium from murine bone marrow-derived macrophages were utilized as positive controls for the expression of active CASP-1 and mature IL-1β, as previously described^54^.

### Immunoblotting

Cell lysates or culture supernatants from murine bone marrow-derived macrophages (30 μg per well; positive control), amniotic fluid (20 µg total protein per well), fetal membrane lysates (75 μg per well), and uterine tissue lysates (75 μg per well) were subjected to electrophoresis in sulfate-polyacrylamide gels (Cat# NP0336BOX; Invitrogen). The proteins were transferred onto nitrocellulose membranes (Cat# 1620145; Bio-Rad, Hercules, CA, USA), which were then submerged in 5% blotting-graded blocking solution (Cat# 1706404; Bio-Rad) for 30 min at room temperature. Next, the membranes were probed overnight at 4°C with anti-mouse CASP-1 (Cat# 14-9832-82; Thermo-Fisher Scientific) or anti-mouse mature IL-1β (Cat# 63124S; Cell Signaling Technology, Danvers, MA, USA; only for fetal membranes). After incubation with each primary antibody, membranes were incubated with HRP-conjugated anti-rat IgG (Cat# 7077S; Cell Signaling) for CASP-1 or HRP-conjugated anti-heavy chain of rabbit IgG (Cat# HRP-66467; Proteintech, Rosemont, IL, USA) for mature IL-1β for 1 h at room temperature. The ChemiGlow West Chemiluminescence Substrate Kit (Cat# 60-12596-00; ProteinSimple, San Jose, CA, USA) was used to detect chemiluminescence signals, and corresponding images were acquired using the ChemiDoc Imaging System (Bio-Rad). The membranes that were loaded with fetal membrane and uterine tissue lysates were re-probed for 1 h at room temperature with a mouse anti-β-actin (ACTB) monoclonal antibody (Cat# A5441, Sigma-Aldrich). Then, the membranes were incubated with HRP-conjugated anti-mouse IgG (Cat# 7076S; Cell Signaling) for 30 min at room temperature. The chemiluminescence signals of the ACTB were detected as shown above. Finally, quantification was performed using ImageJ software as previously reported ^54^. In short, each individual protein band was automatically quantified on the blot images. The internal control, β-actin, was used to normalize the target protein expression in each fetal membrane and uterine tissue sample for relative quantification.

### Immunoprecipitation of mature IL-1β

Immunoprecipitation of cleaved IL-1β from uterine tissue lysates and amniotic fluid was performed using the Pierce™ Classic IP Kit (Cat# 26146; Thermo Fisher Scientific) following the manufacturer’s instructions. Amniotic fluid samples (n = 3 per study group) were pooled. Culture supernatants from murine bone marrow-derived macrophages were utilized as positive controls. Each uterine tissue lysate (1 mg protein) or pooled amniotic fluid sample (1 mg protein) was pre-cleared using the control agarose resin and incubated with rabbit anti-mouse mature IL-1β antibody overnight at 4°C to form the immune complex. Next, the immune complex was captured using Pierce Protein A/G Agarose. After several washes to remove non-bound proteins, the immune complex was eluted with sample buffer and subjected to electrophoresis in 4%–12% polyacrylamide gels followed by Western blot transfer, as described above. The blot was then incubated with rabbit anti-mouse mature IL-1β antibody, followed by incubation with an HRP-conjugated anti-heavy chain of rabbit IgG antibody. Finally, chemiluminescence signals were detected with the ChemiGlow West Substrate Kit and images were acquired using the ChemiDoc Chemiluminescence Imaging System, and quantification was performed using ImageJ software.

### Tracking transferred M2 macrophages *in vivo*

Bone marrow from B6 CD45.1 mice was collected, isolated, and differentiated into M2 macrophages, as described above. Pregnant C57BL/6 recipient mice were intravenously injected with 2-5×10^6^ CD45.1^+^ donor M2 macrophages at 13.5 and 14.5 dpc followed by intra-amniotic injection of HMGB1 on 14.5 dpc, as described above. Mice were euthanized at 2, 6, or 12 h after intra-amniotic injection to collect the maternal blood, maternal lung, uterus, decidua, and placenta as well as the fetal membrane, fetal intestine, fetal liver, fetal liver, and amniotic fluid. Whole maternal blood (100 µL) was used directly for immunophenotyping following the procedure described below. For maternal and fetal tissues, leukocytes were isolated by following protocols adapted from^86^. Briefly, maternal and fetal tissues were gently minced using fine scissors and enzymatically digested with StemPro Accutase Cell Dissociation Reagent (Cat# A1110501, Thermo-Fisher) for 30 min at 37°C. Cells were filtered using a 100 μm cell strainer (Cat# 22-363-549, Fisher Scientific, Fair Lawn, NY, USA) followed by washing with PBS prior to immunophenotyping. Immediately after isolation of leukocytes, cell pellets were re-suspended in FACS buffer and pre-incubated with rat anti-mouse CD16/CD32 Fc Block™ (Cat# 553142, clone 2.4G2; BD Bioscience, Franklin Lakes, NJ, USA) for 10 min on ice and subsequently incubated with specific fluorochrome-conjugated monoclonal anti-mouse antibodies shown in Supplementary Table 3. Cells were acquired using the BD LSRFortessa flow cytometer (BD Biosciences) with FACSDiva 9.0 software (BD Biosciences). CountBright absolute counting beads (Thermo Fisher Scientific) were added prior to acquisition. The analysis was performed and the figures were created by using FlowJo software v10 (FlowJo, Ashland, OR, USA).

### Sampling and gene expressing profiling of neonates

Neonates born to dams that received M2 macrophages or PBS followed by HMGB1 were sacrificed at 14 days of life, and the lung and intestine (cecum, colon, and small intestine) were preserved in RNAlater Stabilization Solution. Gene expression analysis was performed using the methods described above (see “RNA isolation, cDNA synthesis, and qPCR analysis”).

### Neonatal sampling for gut microbiome analysis

Dams underwent treatment with M2 macrophages alone, or were injected with HMGB1 and treated with vehicle or M2 macrophages, as described above. A group of non-injected dams (control) was also included. There was no group-level separation of these dams; each was housed alone after receiving their respective injections. After delivery, dams were housed with their neonates. Surviving neonates were euthanized on postnatal day 14-16. Samples from the neonatal small intestine, cecum, and colon as well as environmental controls were obtained using sterile swabs (FLOQSwabs, Cat# 553C Copan Diagnostics, Murieta, CA, USA) under aseptic conditions. Each experimental group included samples obtained from neonates born to three different dams. All swab samples collected for 16S rRNA gene sequencing were stored at -80°C until DNA extractions were performed.

#### DNA extraction

Genomic DNA was extracted from swabs of the small intestine, cecum, and colon (n = 35 each) and negative blank DNA extraction kit controls using the DNeasy PowerLyzer Powersoil kit (Qiagen, Germantown, MD, USA), with minor modifications to the manufacturer’s protocol as previously described^87–89^. All swabs were randomized across extraction runs. The purified DNA was stored at -20°C.

#### 16s rRNA Gene Sequencing

The V4 region of the 16S rRNA gene was amplified and sequenced via the dual indexing strategy developed by Kozich et al.^90^ as previously described^87–89,91^, with the exception that library builds were performed using 32 cycles of PCR prior to the equimolar pooling of all sample libraries for multiplex sequencing. 16S rRNA gene sequences were clustered into amplicon sequence variants (ASVs) defined by 100% sequence similarity using DADA2 version 1.12^92^ in R version 4.2.2^93^ as previously described^94^, with the exception that forward and reverse reads were truncated at 240 and 215 bases, respectively. Sequences were then classified using the silva_nr_v132_train_set database with a minimum bootstrap value of 80%, and sequences that were derived from Archaea, chloroplast, Eukaryota, or mitochondria were removed.

Potential background DNA contaminant ASVs were identified using the R package decontam version 1.18.0^95^ and the “IsContaminant” function. One of the 82 ASVs identified as a contaminant [Lachnospiraceae (ASV-2)] was present in > 90% of both biological and control samples, however, it had a greater mean relative abundance in biological (12.95%) than in control samples (1.77%) and was therefore not removed from the dataset. The two taxa with the highest mean prevalence and abundance in control samples that were removed from the dataset were Burkholderiaceae (ASV-42) (41.7% and 5.3%) and *Pseudomonas* (ASV-97) (50.0% and 3.8%, respectively). The mean relative abundances of these two ASVs across all biological samples were 0.0003% and 0.006%, respectively. Other ASVs removed from the dataset included taxa that have previously been reported to be contaminants in 16S rRNA gene sequencing studies (e.g., *Acinetobacter*, *Cloacibacterium*, *Corynebacterium*, *Cutibacterium*, *Escherichia*, *Halomonas*, *Pseudomonas*, *Ralstonia*, *Sphingomonas*, *Staphylococcus*, *Stenotrophomonas*, and *Xanthomonas*)^91,96–99^.

After removing contaminant ASVs, all samples were normalized to 8,715 reads using the “rarefy_even_depth” function in phyloseq 1.42.0^100^, resulting in 20 bacterial profiles with Good’s coverage^101^ values ≥ 99.68% for each of the three sample types. The final dataset contained a total of 411 ASVs. No cross-tissue comparisons were conducted.

#### Beta diversity

Beta diversity of bacterial profiles was characterized using the Bray-Curtis similarity index. Principal coordinate analysis (PCoA) plots were used to visualize the Bray-Curtis similarity of the sample profiles. Using nonparametric multivariate analysis of variance (NPMANOVA) as implemented in vegan 2.6.4^102^, differences in bacterial community structure across treatments were evaluated for each tissue type.

#### Differential abundance of bacterial species

Differential abundance of the 25 most prevalent ASVs between treatment groups was independently evaluated for the small intestine, cecum, and colon datasets using two-tailed Mann-Whitney U-tests with Holm’s correction for multiple comparisons as implemented in R version 4.2.2. Adjusted p-values < 0.1 were considered significant.

### Neonatal bacterial challenge

Group B *Streptococcus* (GBS, CNCTC 10/84, serotype V, sequence type 26) was originally isolated from a neonate with sepsis. From an overnight culture, a sub-culture was placed with fresh Tryptic Soy Broth (Cat# 211825; BD Bioscience, Franklin Lakes, NJ, USA) and grown to the logarithmic phase (OD600 0.8-0.9). Additional dilution was performed using sterile PBS to reach working concentrations. Thriving term neonates (14-16 days old) born to dams that did not receive any treatment (control) and dams injected solely with HMGB1, PBS (vehicle for M2 macrophages) followed by HMGB1, or M2 macrophages followed by HMGB1 were intraperitoneally injected with 2×10^6^ colony-forming units of GBS in 200 µL sterile PBS. Survivability, body weight, and rectal temperature of the challenged neonates were checked at 4-6 h intervals throughout the daytime for 5 days.

## QUANTIFICATION AND STATISTICAL ANALYSIS

Statistical analyses were performed using GraphPad Prism (v9.5.0; GraphPad, San Diego, CA, US) as indicated in each figure caption. Flow cytometry analysis was performed using FlowJo software v10. Protein expression quantification was performed using ImageJ software. To determine gene expression levels from qPCR arrays, -ΔC_T_ values were calculated using averaged reference genes (*Actb, Gapdh, Gusb*, and *Hsp90ab1*) within each sample. Heatmaps were created to represent the Z-scores. Single-cell RNA-sequencing and microbiome data analysis were performed using R (v.4.2.2; https://www.r-project.org/), as described above. A p-value ≤ 0.05 or adjusted p-value (q-value) ≤ 0.1 was considered statistically significant.

## Supplemental Items

**Document S1.** Figures S1-S11 and Tables S2-S3

**Table S1.** Excel file containing additional data too large to fit in a PDF, related to Figure 1

## Supplemental Item Legends

**Figure S1.** Gene Ontology of macrophage populations at the maternal-fetal interface, related to Figure 1.

**Figure S2.** Intra-amniotic injection of HMGB1 does not significantly alter amniotic fluid concentrations of total IL-1β and TNF, related to Figure 3.

**Figure S3.** M2 macrophages do not significantly alter HMGB1-induced inflammasome activation in the uterine tissues.

**Figure S4.** Neither HMGB1 nor M2 macrophage treatment induce changes in uterine gene expression.

**Figure S5.** M2 macrophages dampen inflammatory gene expression in the decidua.

**Figure S6.** Adoptively transferred M2 macrophages accumulate at the maternal-fetal interface but do not reach fetal organs.

**Figure S7.** M2 macrophages ameliorate the HMGB1-induced inflammatory response in the placenta.

**Figure S8.** HMGB1 regulates gene expression in the fetal intestine, related to Figure 6.

**Figure S9.** M2 macrophages change the neonatal gut microbiome after in utero exposure to HMGB1, related to Figure 6.

**Figure S10.** M2 macrophages modulate microbiome dysbiosis in each compartment of the neonatal gut, related to Figure 6.

**Figure S11.** M2 macrophages reduce neonatal hypothermia upon bacterial challenge, related to Figure 7.

**Table S1.** Marker genes used to distinguish macrophage clusters M1-M7, related to Figure 1

**Table S2.** TaqMan assays utilized for qPCR.

**Table S3.** Antibodies utilized for flow cytometry.

