## Supplementary material for "Homeostatic macrophages prevent preterm birth and improve neonatal outcomes by mitigating *in utero* sterile inflammation": Key Resource Table

**Key resources table**

| REAGENT or RESOURCE | SOURCE | IDENTIFIER |
| --- | --- | --- |
| Antibodies | | |
| Monoclonal rat anti-mouse F4/80 antibody; clone T45-2342 | BD Biosciences | Cat# 565613, RRID:AB_2734770 |
| Monoclonal mouse anti-mouse CD45.1 antibody; clone A20 | BD Biosciences | Cat# 553775, RRID:AB_395043 |
| Monoclonal mouse anti-mouse CD45.2; clone 104 | BD Biosciences | Cat# 560694, RRID:AB_1727492 |
| Rat IgG2a, κ isotype control; clone R35-95 | BD Biosciences | Cat# 562302, RRID:AB_11154396 |
| Mouse IgG2a, κ isotype control; clone G155-178 | BD Biosciences | Cat# 553456, RRID:AB_479604 |
| Mouse IgG2a, κ isotype control; clone G155-178 | BD Biosciences | Cat# 557751, RRID:AB_396857 |
| Rat anti-mouse CD16/CD32 Fc Block antibody; clone 2.4G2 | BD Biosciences | Cat# 553142, RRID: RRID:AB_394657 |
| Monoclonal rat anti-mouse CASP-1 antibody; clone 5B10 | Thermo-Fisher Scientific | Cat# 14-9832-82, RRID:AB_2016691 |
| Monoclonal rabbit anti-mouse mature IL-1β antibody; clone E7V2A | Cell Signaling Technology | Cat# 63124S |
| Rat IgG antibody | Cell Signaling Technology | Cat# 7077S |
| Rabbit heavy chain IgG antibody; clone 1C9A5 | Proteintech | Cat# HRP-66467, RRID:AB_2883840 |
| Monoclonal mouse anti-β-actin antibody; clone AC-15 | Sigma-Aldrich | Cat# A5441, RRID:AB_476744 |
| Mouse IgG antibody | Cell Signaling Technology | Cat# 7076S |
| Bacterial and virus strains | | |
| Group B *Streptococcus* (GBS, CNCTC 10/84, serotype V, sequence type 26) isolated from a neonate with sepsis | This manuscript |  |
| Biological samples |  |  |
| Chemicals, peptides, and recombinant proteins | | |
| Recombinant HMGB1 | BioLegend | Cat# 557804 |
| Recombinant mouse macrophage colony-stimulating factor (M-CSF) | BioLegend | Cat# 576402 |
| Recombinant mouse IL-4 | BioLegend | Cat# 574302 |
| Recombinant mouse IL-13 | BioLegend | Cat# 575902 |
| Protease inhibitor cocktail | Roche Applied Sciences | Cat# 11836170001 |
| Critical commercial assays | | |
| U-PLEX Custom Biomarker Group 1 | Meso Scale Discovery | Cat# K15069M-2 |
| Pierce BCA Protein Assay Kit | Pierce Biotechnology | Cat# 23225 |
| ChemiGlow West Chemiluminescence Substrate Kit | ProteinSimple | Cat# 60-12596-00 |
| Pierce Classic IP kit | Thermo-Fisher | Cat# 26146 |
| Deposited data | | |
| Microbial 16S rRNA sequencing of the neonatal intestine | This manuscript | NCBI Sequence Read Archive, Accession Number PRJNA925285 |
| Single-cell RNA-sequencing data of the human placenta | Pique-Regi, R et al. 2019^44^ | NIH dbGAP, Accession Number, phs001886.v5.p1 |
| Single-cell RNA-sequencing data of the human myometrium | Pique-Regi, R et al. 2022^45^ | NIH dbGAP, Accession Number phs001886.v5.p1 |
| Single-cell RNA-sequencing data of the human placenta | Garcia-Flores, V et al. 2024^46^ | NIH dbGAP, Accession Number phs001886.v5.p1 |
| Experimental models: Cell lines | | |
| Experimental models: Organisms/strains | | |
| C57BL/6 mice | The Jackson Laboratory | Stock# 000664 |
| B6 CD45.1 mice | The Jackson Laboratory | Stock# 002014 |
| Oligonucleotides | | |
| TaqMan assays for RT-qPCR, see Table S3 |  |  |
| Recombinant DNA | | |
| Software and algorithms | | |
| GraphPad Prism v9.5.0 | GraphPad |  |
| ImageJ software | NIH |  |
| R package 4.2.2 | R-project |  |
| kallisto | Bray, NL et al. 2016^76^ |  |
| bustools | Melsted, P et al. 2021^77^ |  |
| STAR aligner | Dobin, A et al. 2013^78^ |  |
| souporcell | Heaton, H et al. 2020^79^ |  |
| demuxlet | Kang, HM et al. 2018^80^ |  |
| Seurat | Stuart, T et al. 2019^81^; Hafemeister, C and Satija, R 2019^82^ |  |
| Harmony | Korsunsky, I et al. 2019^83^ |  |
| DADA2 | Callahan, BJ et al. 2016^92^ |  |
| decontam | Davis, NM et al. 2018^95^ |  |
| phyloseq | McMurdie, PJ et al. 2013^100^ |  |
| vegan | Dixon, P 2003^102^ |  |
| Cell Ranger v4.0.0 | 10x Genomics |  |
| FACSDiva v9.0 | BD Biosciences |  |
| FlowJo v10 | FlowJo |  |
| Other | | |
| QIAshredders | Qiagen | Cat# 79656 |
| RNAlater stabilization solution | Invitrogen | Cat# AM7021 |
| RNase-free DNase | Qiagen | Cat# 79254 |
| RNeasy Mini Kit | Qiagen | Cat# 74104 |
| SuperScript IV VILO master mix | Invitrogen | Cat# 11756050 |
| DNeasy PowerLyzer Powersoil kit | Qiagen | Cat# 47014 |
