## Supplementary Figures 1-11 and Tables 2-3 for "Homeostatic macrophages prevent preterm birth and improve neonatal outcomes by mitigating *in utero* sterile inflammation"

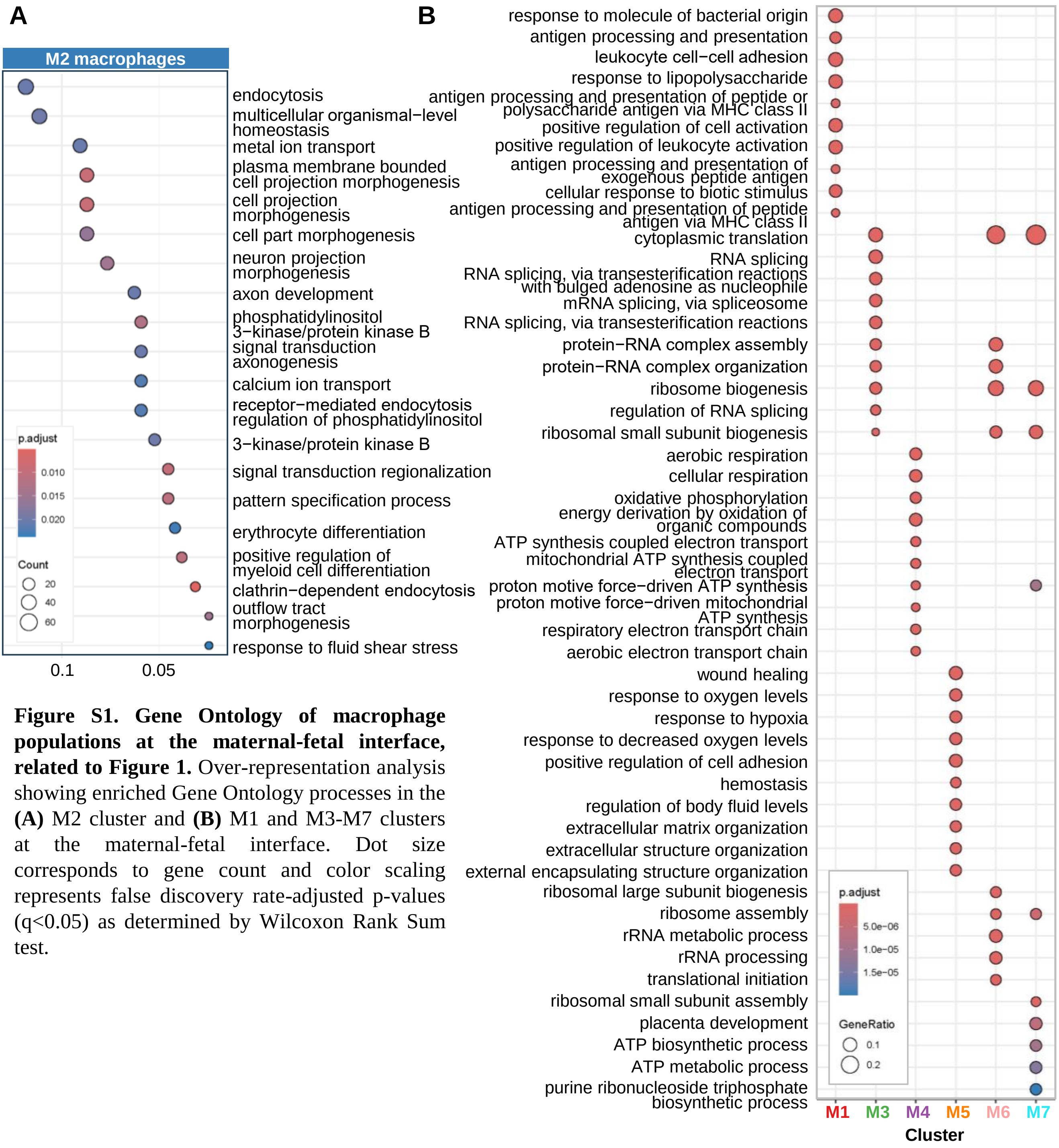


**Figure S1. Gene Ontology of macrophage populations at the maternal-fetal interface, related to Figure 1.** Over-representation analysis showing enriched Gene Ontology processes in the **(A)** M2 cluster and **(B)** M1 and M3-M7 clusters at the maternal-fetal interface. Dot size corresponds to gene count and color scaling represents false discovery rate-adjusted p-values (q<0.05) as determined by Wilcoxon Rank Sum test.

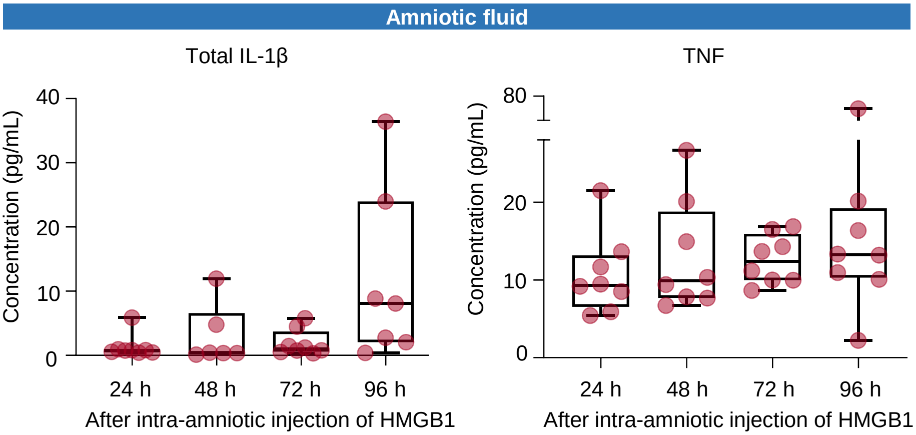


**Figure S2. Intra-amniotic injection of HMGB1 does not significantly alter amniotic fluid concentrations of total IL-1β and TNF, related to Figure 3.** Concentrations of total IL-1β and TNF in the amniotic fluid of HMGB1-injected dams collected at 24, 48, 72, or 96 h post-injection (n = 6-8 dams per time point). Statistical analysis was performed using the Kruskal-Wallis test followed by two-stage linear step-up procedure of Benjamini, Krieger, and Yekutieli post-hoc test. Data are shown as box plots where midlines indicate medians, boxes denote interquartile ranges, and whiskers indicate the minimum/maximum range.

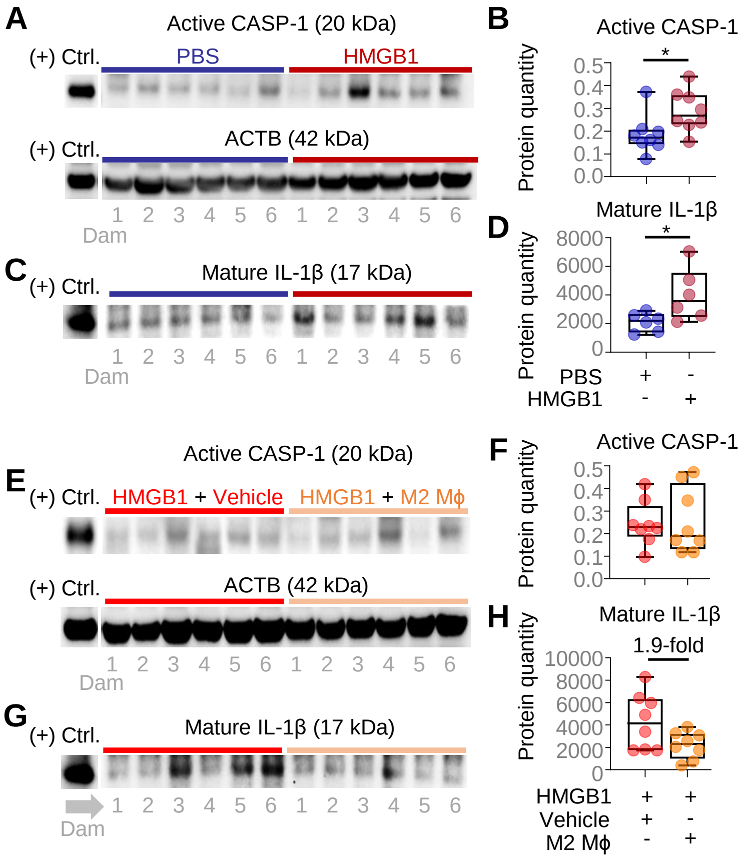


**Figure S3. M2 macrophages do not significantly alter HMGB1-induced inflammasome activation in the uterine tissues.** Dams received intra-amniotic injection of PBS (control) or HMGB1 on 14.5 days *post coitum* (dpc). The uterus was collected at 72 h post-HMGB1 injection for immunoblotting. **(A)** Immunoblotting of active caspase-1 (CASP-1) and β-actin (ACTB) expression in the uterine tissues from PBS- or HMGB1-injected dams. Representative immunoblot images depict 6 samples per group in each gel. **(B)** Protein quantification of active CASP-1 (normalized by ACTB) in the uterus of PBS- or HMGB1-injected dams (n = 8 per group). **(C)** Immunoblotting of mature IL-1β expression in the uterus following immunoprecipitation with anti-IL-1β antibody. The representative immunoblot image depicts 6 samples per group. **(D)** Protein quantification of mature IL-1β in the uterus of PBS- or HMGB1-injected dams (n = 6 per group). M2 macrophages (M2 MΦ) or PBS (Vehicle) were intravenously administered to dams on 13.5 dpc and 14.5 dpc followed by intra-amniotic injection of HMGB1 on 14.5 dpc. The uterus was collected at 72 h post-HMGB1 injection for immunoblotting. **(E)** Immunoblotting of active CASP-1 and ACTB expression in the uterine tissues from HMGB1+Vehicle or HMGB1+M2 MΦ dams. Representative immunoblot images depict 6 samples per group in each gel. **(F)** Protein quantification of active CASP-1 (normalized by ACTB) in the uterus of HMGB1+Vehicle or HMGB1+M2 MΦ dams (n = 8 per group). **(G)** Immunoblotting of mature IL-1β expression in the uterus following immunoprecipitation with anti-IL-1β antibody. The representative immunoblot image depicts 6 samples per group. **(H)** Protein quantification of mature IL-1β in the uterus of HMGB1+Vehicle or HMGB1+M2 MΦ dams (n = 8 per group). P-values were determined using the two-tailed Mann-Whitney U-test. Data are shown as boxplots where midlines indicate medians, boxes denote interquartile ranges, and whiskers indicate the minimum/maximum range. *$p < 0.05$.

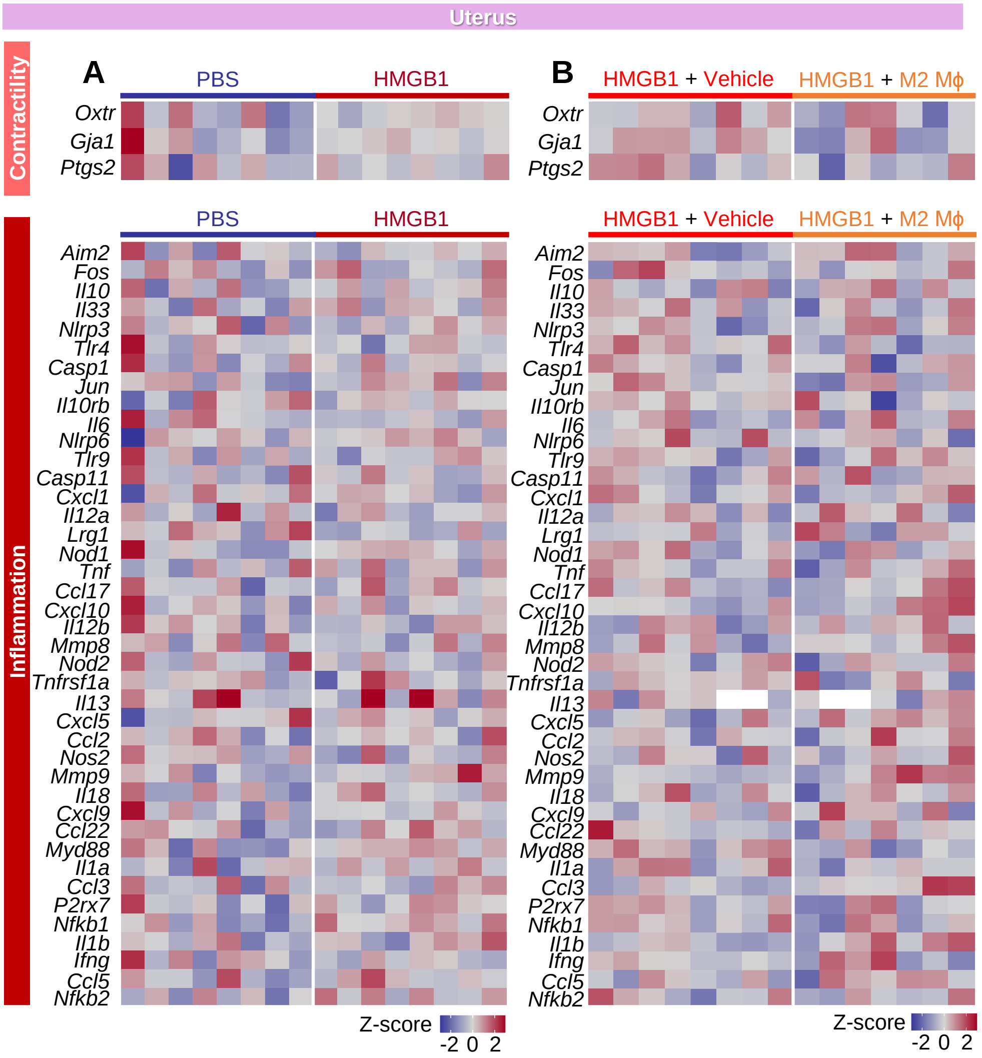


**Figure S4. Neither HMGB1 nor M2 macrophage treatment induce changes in uterine gene expression. (A)** Dams received intra-amniotic injection of PBS (control) or HMGB1 on 14.5 days *post coitum* (dpc) followed by tissue collection at 72 h post-injection to determine contractility-associated and inflammatory gene expression. Heatmaps representing gene expression in the uterus of PBS- or HMGB1-injected dams (n = 8 per group). **(B)** M2 macrophages (M2 MΦ) or PBS (Vehicle) were intravenously administered to dams on 13.5 dpc and 14.5 dpc followed by intra-amniotic injection of HMGB1 on 14.5 dpc. Tissues were collected at 72 h post-HMGB1 injection to determine contractility-associated and inflammatory gene expression. Heatmaps representing gene expression in the uterus of HMGB1+Vehicle (n = 8) or HMGB1+M2 MΦ (n = 7) dams.

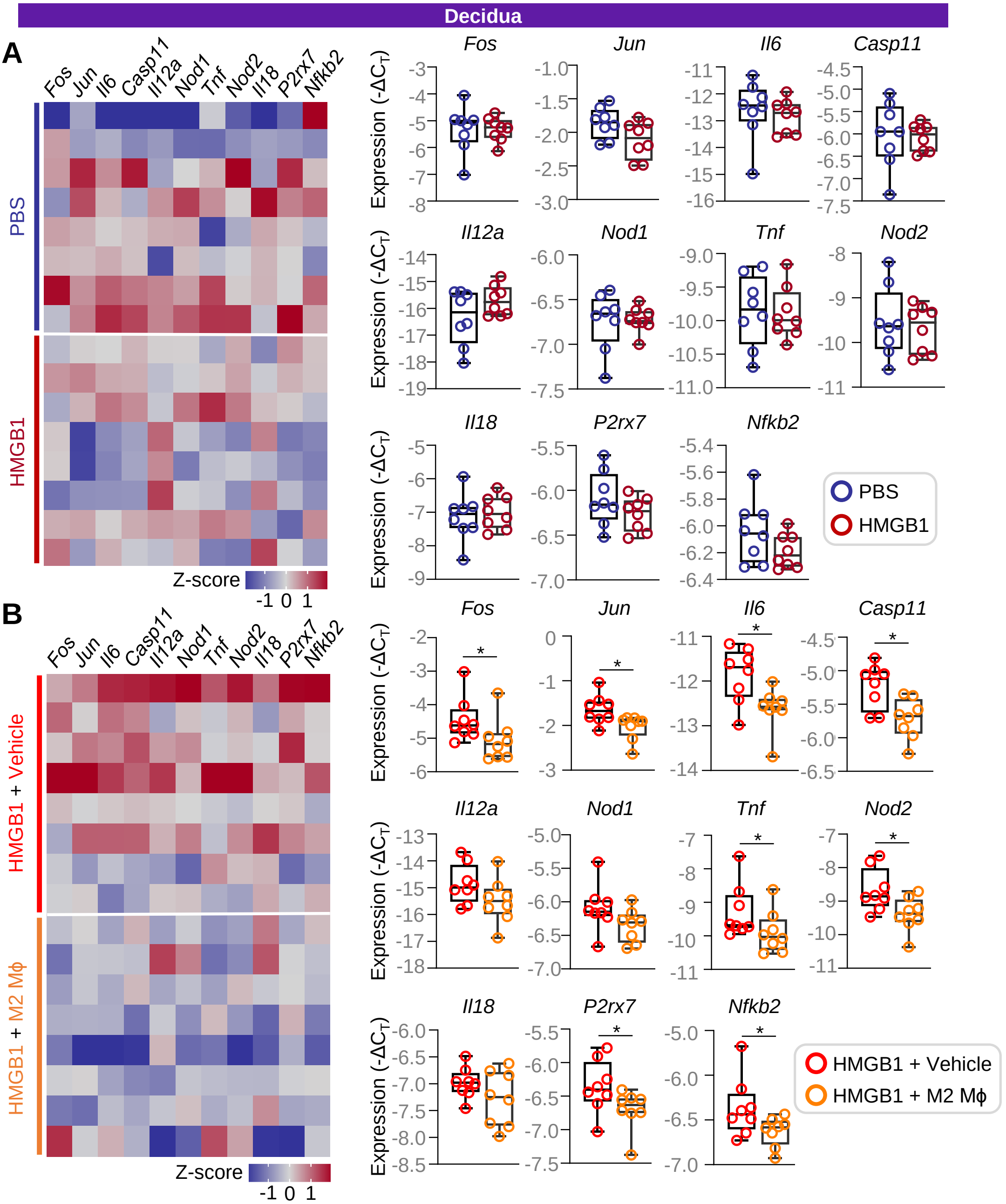


**Figure S5. M2 macrophages dampen inflammatory gene expression in the decidua. (A)** Dams received intra-amniotic injection of PBS (control) or HMGB1 on 14.5 days *post coitum* (dpc). Tissue collection was performed at 72 h post-injection to collect the decidua for gene expression profiling. Representative heatmaps displaying the expression of key inflammatory genes in the decidua of PBS- or HMGB1-injected dams. Expression of *Fos*, *Jun*, *Il6*, *Casp11*, *Il12a*, *Nod1*, *Tnf*, *Nod2*, *Il18*, *P2rx7*, and *Nfkb2* in the decidua of PBS- or HMGB1-injected dams (n = 8 per group). **(B)** M2 macrophages (M2 MΦ) or PBS (Vehicle) were intravenously administered to dams on 13.5 dpc and 14.5 dpc followed by intra-amniotic injection of HMGB1 on 14.5 dpc. The decidua was collected at 72 h post-HMGB1 injection to determine gene expression. Representative heatmaps displaying the expression of key inflammatory genes in the decidua of HMGB1+Vehicle or HMGB1+M2 MΦ dams. Expression of *Fos*, *Jun*, *Il6*, *Casp11*, *Il12a*, *Nod1*, *Tnf*, *Nod2*, *Il18*, *P2rx7*, and *Nfkb2* in the decidua of HMGB1+Vehicle or HMGB1+M2 MΦ dams (n = 8 per group). Data are shown as box plots where midlines indicate medians, boxes denote interquartile ranges, and whiskers indicate the minimum/maximum range. P-values were determined using the two-tailed Mann-Whitney U-test. *$p < 0.05$.

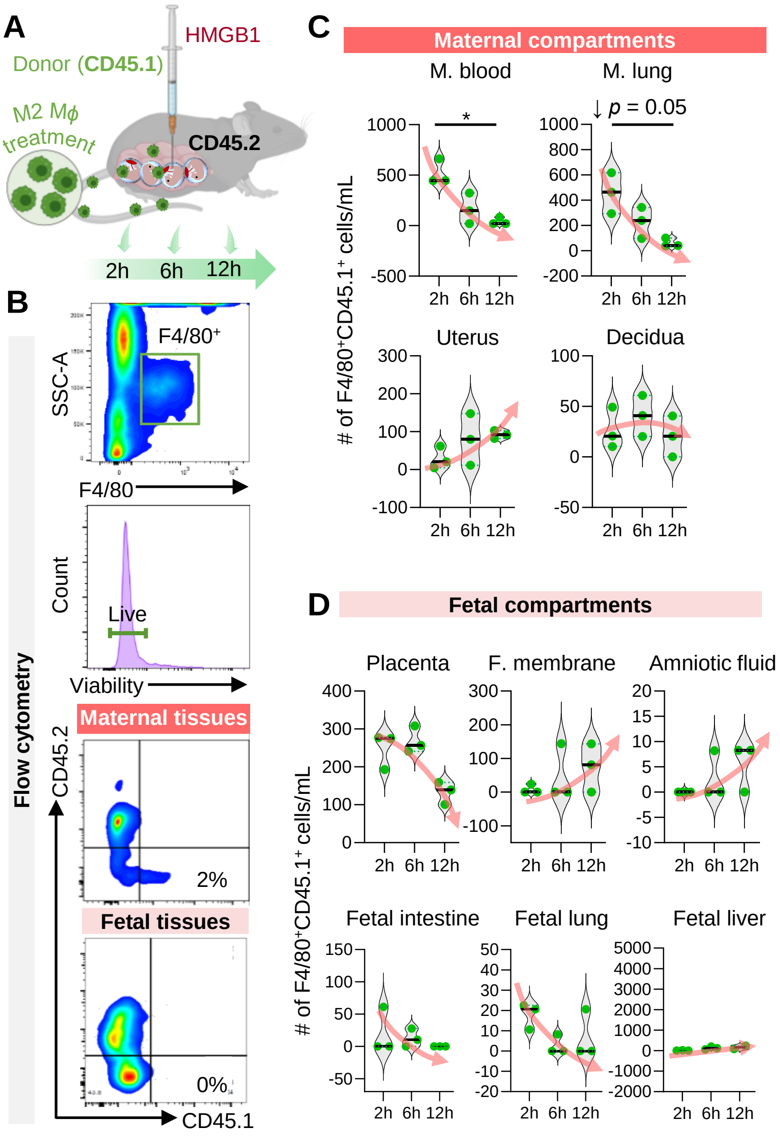


**Figure S6. Adoptively transferred M2 macrophages accumulate at the maternal-fetal interface but do not reach fetal organs. (A)** Bone marrow-derived cells were collected from donor CD45.1+ mice, differentiated, and polarized to M2 macrophages (M2 MΦ) *in vitro*. Donor M2 MΦ were intravenously administered on 13.5 days *post coitum* (dpc) and 14.5 dpc to recipient CD45.2+ dams followed by intra-amniotic injection of HMGB1 on 14.5 dpc. Maternal and fetal tissues were collected at 2, 6, or 12 h post-HMGB1 injection (n = 2-3 dams per time point) to track donor M2 MΦ infiltration. **(B)** Representative flow cytometry gating strategy to detect viable CD45.1+CD45.2- donor macrophages in maternal and fetal tissues. **(C)** Quantification of CD45.1+CD45.2- donor macrophages in the maternal blood, maternal lung, uterus, and decidua from recipient dams at 2, 6, and 12 h post-HMGB1 injection. **(D)** Quantification of CD45.1+CD45.2- donor macrophages in the placenta, fetal membranes, amniotic fluid, fetal intestine, fetal lung, and fetal liver from recipient dams at 2, 6, and 12 h post-HMGB1 injection. Data are presented as violin plots where the midline represents the median, dotted lines represent the interquartile range (IQR), and whiskers represent the minimum/maximum range. Statistical analysis was performed by using the Kruskal-Wallis test followed by Dunn’s post-hoc test. $^*p<0.05$

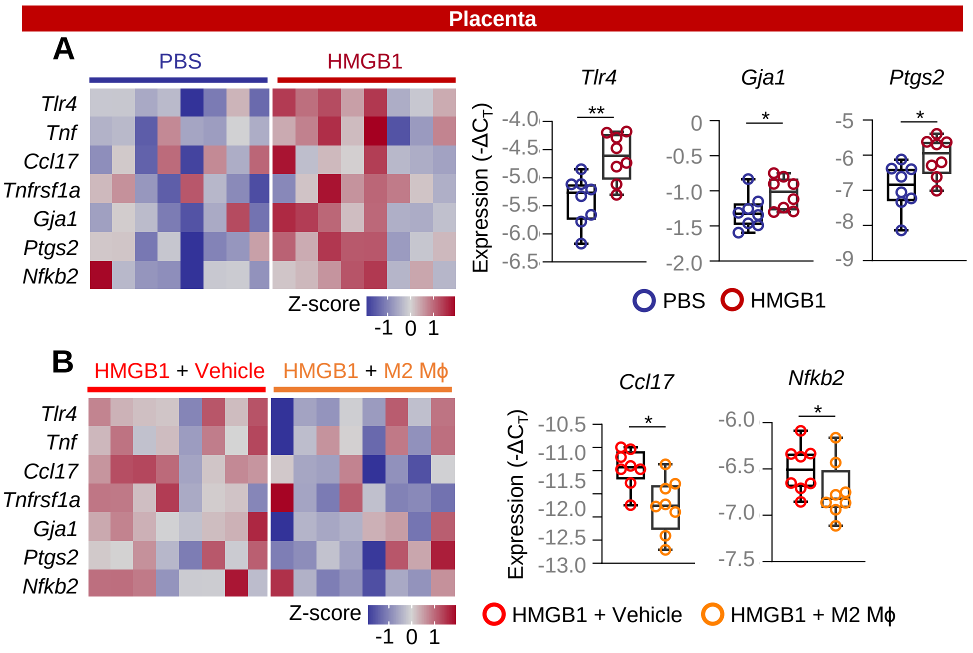


**Figure S7. M2 macrophages ameliorate the HMGB1-induced inflammatory response in the placenta. (A)** Dams received intra-amniotic injection of PBS (control) or HMGB1 on 14.5 days *post coitum* (dpc). Tissue collection was performed at 72 h post-injection to collect the placenta for gene expression profiling. Representative heatmaps displaying the expression of inflammatory or contractility-associated genes in the placenta of PBS- or HMGB1-injected dams. Expression of *Tlr4*, *Gja1*, and *Ptgs2* in the placenta of PBS- or HMGB1-injected dams (n = 8 per group). **(B)** M2 macrophages (M2 MΦ) or PBS (Vehicle) were intravenously administered to dams on 13.5 dpc and 14.5 dpc followed by intra-amniotic injection of HMGB1 on 14.5 dpc. The placenta was collected at 72 h post-HMGB1 injection for gene expression profiling. Representative heatmaps displaying the expression of inflammatory or contractility-associated genes in the placenta of HMGB1+Vehicle or HMGB1+M2 MΦ dams. Expression of *Ccl17* and *Nfkb2* in the placenta of HMGB1+Vehicle or HMGB1+M2 MΦ dams (n = 8 per group). Data are shown as box plots where midlines indicate medians, boxes denote interquartile ranges, and whiskers indicate the minimum/maximum range. P-values were determined using the two-tailed Mann-Whitney U-test. $^*p < 0.05$; $^{**}p < 0.01$.

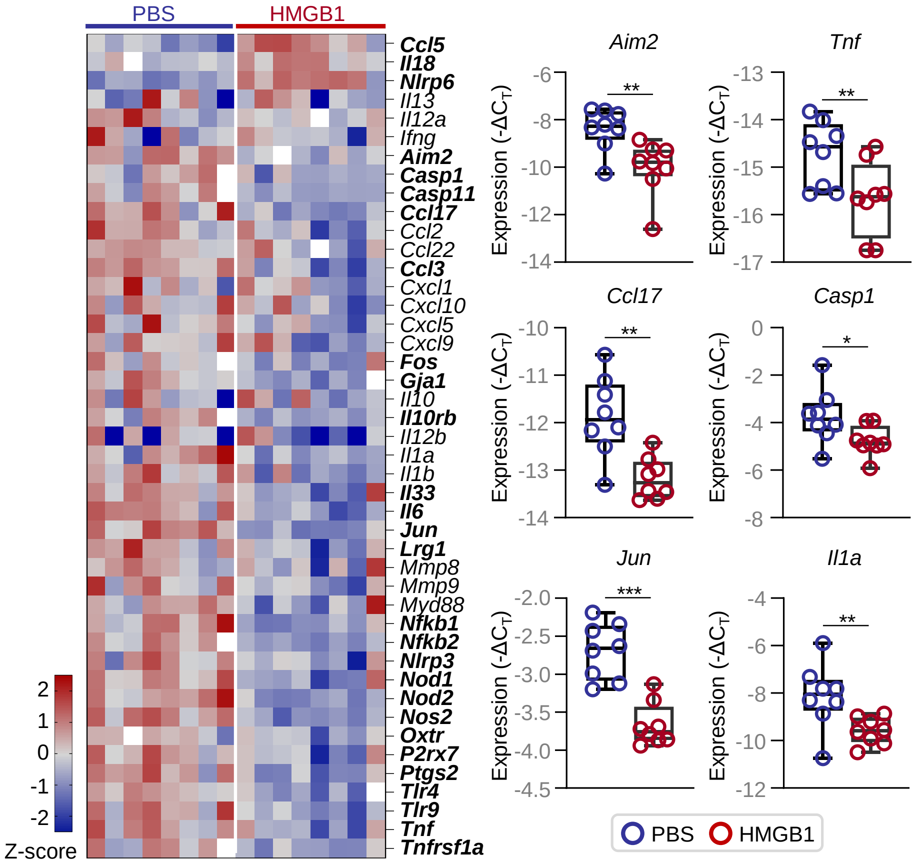


**Figure S8. HMGB1 regulates gene expression in the fetal intestine, related to Figure 6.** Dams received intra-amniotic injection of PBS (control) or HMGB1 on 14.5 days *post coitum* (dpc). Tissue collection was performed at 72 h post-injection to collect fetal intestines and determine gene expression as shown in the representative heatmap. Genes in bold font are statistically significant between groups. Expression of *Aim2, Tnf, Ccl17, Casp1, Jun,* and *Il1a* in the fetal intestine (n = 8 per group). Data are shown as boxplots where midlines indicate medians, boxes denote interquartile ranges, and whiskers indicate the minimum/maximum range. P-values were determined using the two-tailed Mann-Whitney U-test. *p < 0.05; **p < 0.01; ***p < 0.001.

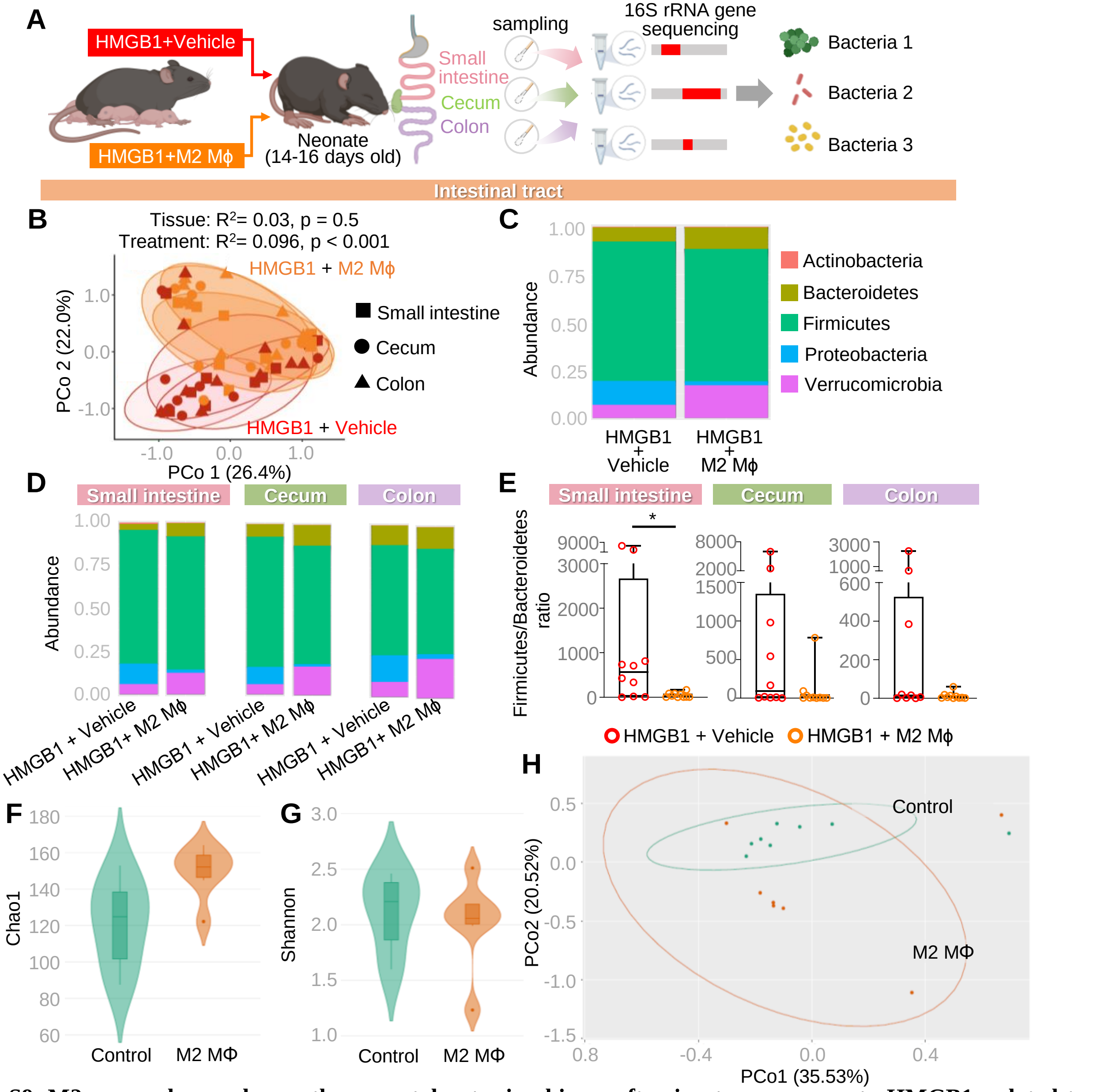


**Figure S9. M2 macrophages change the neonatal gut microbiome after *in utero* exposure to HMGB1, related to Figure 6. (A)** M2 macrophages (M2 MΦ) or PBS (Vehicle) were intravenously administered to dams on 13.5 days *post coitum* (dpc) and 14.5 dpc followed by intra-amniotic injection of HMGB1 on 14.5 dpc. Neonates were monitored until 14-16 days of age, and the small intestine, cecum, and colon were collected for microbial analysis using 16S rRNA gene sequencing (n = 10 neonates per group). **(B)** Principal coordinates analysis (PCoA) illustrating variation in the 16S rRNA gene profiles according to tissue or treatment (n = 10 per group). Similarities in the 16S rRNA gene profiles were characterized using the Bray-Curtis similarity index. **(D-E)** Differences in abundance of major microbial phyla and change in the Firmicutes/Bacteroidetes ratio in the neonatal small intestine, neonatal cecum, and neonatal colon. Data are shown as box plots where midlines indicate medians, boxes denote interquartile ranges, and whiskers indicate the minimum/maximum range. P-values were determined using two-tailed Mann-Whitney U-tests (for comparing abundance of individual taxa). Alpha diversity metrics **(F)** Chao1 index (i.e., richness) and **(G)** Shannon index (i.e., heterogeneity) for the microbiota composition of intestinal tissues (combined small intestine, cecum, and colon) from neonates born to non-injected dams (control) or dams that received M2 macrophages (M2 MΦ) alone (n = 7-8 neonates per group). **(H)** Principal coordinates analysis (PCoA) illustrating the Bray-Curtis structural variation in the 16S rRNA gene profiles between groups. Ellipses were drawn for each group at the 95% confidence interval. *$p < 0.05$.

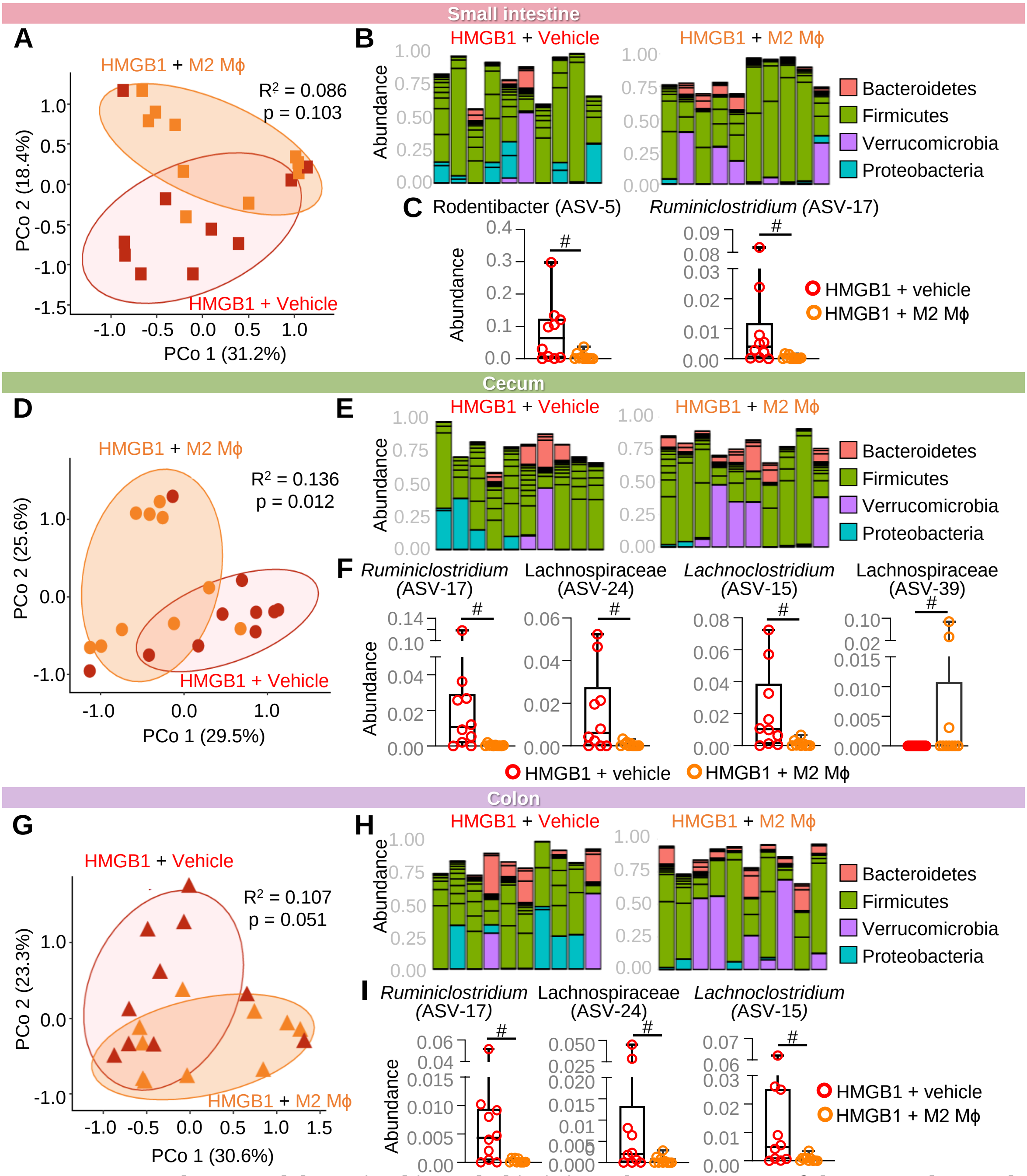


**Figure S10. M2 macrophages modulate microbiome dysbiosis in each compartment of the neonatal gut, related to Figure 6.** Principal coordinates analysis (PCoA) illustrating variation in the 16S rRNA gene profiles among the **(A)** small intestine, **(D)** cecum, and **(G)** colon (n=10 neonates per group). 16S rRNA gene profiles were characterized using the Bray-Curtis similarity index. Bar plots showing taxonomic classifications of 20 bacterial taxa with highest relative abundance across all **(B)** small intestine, **(E)** cecum, and **(H)** colon samples. Multiple taxa with the same bacterial taxonomic classification within a same sample are indicated by bars of the same color. Changes in abundance of **(C)** *Rodentibacter* (ASV-5) and *Ruminiclostridium* (ASV-17) in the neonatal small intestine, **(F)** *Ruminiclostridium* (ASV-17), *Lachnospiraceae* (ASV-24), *Lachnoclostridium* (ASV-15), and *Lachnospiraceae* (ASV-39) in the neonatal cecum, and **(I)** *Ruminiclostridium* (ASV-17), *Lachnospiraceae* (ASV-24), and *Lachnoclostridium* (ASV-15) in the neonatal colon. Data are shown as box plots where midlines indicate medians, boxes denote interquartile ranges, and whiskers indicate the minimum/maximum range. P-values were determined using two-tailed Mann-Whitney U-tests with Holm's correction for multiple comparisons. #q < 0.1.

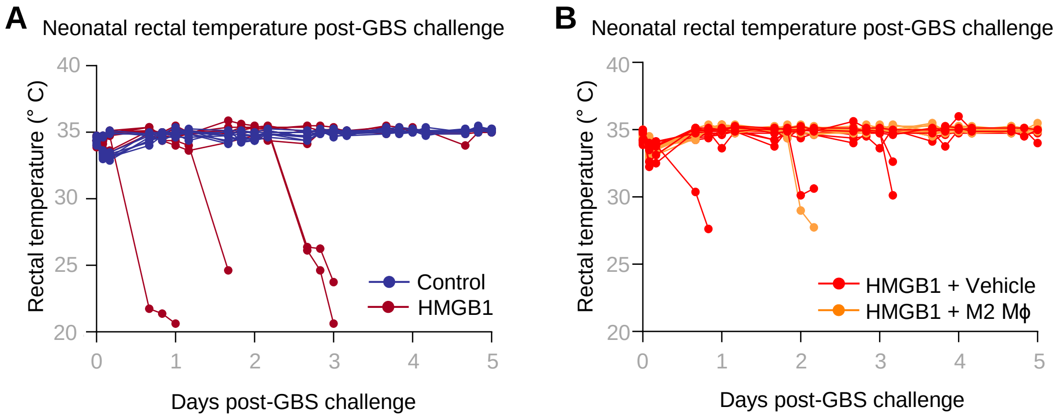


**Figure S11. M2 macrophages reduce neonatal hypothermia upon bacterial challenge, related to Figure 7. (A)** Rectal temperature of group B *Streptococcus* (GBS)-challenged neonates born to dams without treatment or with intra-amniotic HMGB1 over the five days post-challenge (n = 10 per group). **(B)** Rectal temperature of GBS-challenged neonates born to HMGB1+Vehicle or HMGB1+M2 MΦ dams over the five days post-challenge (n = 12 per group).

**Table S2.** TaqMan assays utilized for RT-qPCR

| **Name** | **Symbol** | **Assay ID** |
|---|---|---|
| Actin, beta | *Actb* | Mm04394036_g1 |
| Glucuronidase, beta | *Gusb* | Mm01197698_m1 |
| Glyceraldehyde-3-phosphate dehydrogenase | *Gapdh* | Mm99999915_g1 |
| Heat shock protein 90 alpha (cytosolic), class B member 1 | *Hsp90ab1* | Mm00833431_g1 |
| Absent In Melanoma 2 | *Aim2* | Mm01295719_m1 |
| Arginase 1 | *Arg1* | Mm00475988_m1 |
| Caspase 1 | *Casp1* | Mm00438023_m1 |
| Caspase 11 (caspase 4) | *Casp11* | Mm00432304_m1 |
| Cluster of differentiation 68 | *Cd68* | Mm03047343_m1 |
| Chemokine (C-C motif) ligand 17 | *Ccl17* | Mm01244826_g1 |
| Chemokine (C-C motif) ligand 2 | *Ccl2* | Mm00441242_m1 |
| Chemokine (C-C motif) ligand 22 | *Ccl22* | Mm00436439_m1 |
| Chemokine (C-C motif) ligand 3 | *Ccl3* | Mm00441259_g1 |
| Chemokine (C-C motif) ligand 5 | *Ccl5* | Mm01302427_m1 |
| chemokine (C-X-C motif) ligand 1 | *Cxcl1* | Mm04207460_m1 |
| chemokine (C-X-C motif) ligand 10 | *Cxcl10* | Mm00445235_m1 |
| chemokine (C-X-C motif) ligand 5 | *Cxcl5* | Mm00436451_g1 |
| chemokine (C-X-C motif) ligand 9 | *Cxcl9* | Mm00434946_m1 |
| FBJ osteosarcoma oncogene | *Fos* | Mm00487425_m1 |
| Gap junction protein, alpha 1 | *Gja1* | Mm00439105_m1 |
| Interferon gamma | *Ifng* | Mm01168134_m1 |
| Insulin-like Growth Factor Receptor-1 | *Igf2* | Mm00439564_m1 |
| Interleukin-10 | *Il10* | Mm01288386_m1 |
| Interleukin-10 receptor beta | *Il10rb* | Mm00434157_m1 |
| Interleukin-12a | *Il12a* | Mm00434169_m1 |
| Interleukin-12b | *Il12b* | Mm01288989_m1 |
| Interleukin-13 | *Il13* | Mm00434204_m1 |
| Interleukin-18 | *Il18* | Mm00434226_m1 |
| Interleukin-1 alpha | *Il1a* | Mm00439620_m1 |
| Interleukin-1 beta | *Il1b* | Mm00434228_m1 |
| Interleukin-33 | *Il33* | Mm00505403_m1 |
| Interleukin-6 | *Il6* | Mm00446190-m1 |
| Leucine-rich α-2 glycoprotein 1 | *Lrg1* | Mm01278767_m1 |
| KDM1 lysine (K)-specific demethylase 6B | *Kdm6b* | Mm01332680_m1 |
| Jun proto-oncogene | *Jun* | Mm07296811_s1 |
| Metabotropic glutamate receptor subtype 5 | *mGluR5* | Mm00690332_m1 |
| Matrix Metallopeptidase 8 | *Mmp8* | Mm00439509_m1 |
| Matrix Metallopeptidase 9 | *Mmp9* | Mm00442991_m1 |

| | | |
|---|---|---|
| Myeloid differentiation primary response gene 88 | *Myd88* | Mm00440338_m1 |
| Nuclear factor of kappa light polypeptide gene enhancer in B cells 1 | *Nfkb1* | Mm00476361_m1 |
| Nuclear factor of kappa light polypeptide gene enhancer in B cells 2 | *Nfkb2* | Mm00479807_m1 |
| NLR family, pyrin domain containing 3 | *Nlrp3* | Mm00840904_m1 |
| NLR family, pyrin domain containing 6 | *Nlrp6* | Mm00460229_m1 |
| Nucleotide-binding oligomerization domain-containing protein 1 | *Nod1* | Mm00805062_m1 |
| Nucleotide-binding oligomerization domain-containing protein 2 | *Nod2* | Mm00467543_m1 |
| Nitric oxide synthase 2, inducible | *Nos2* | Mm00440502_m1 |
| Oxytocin receptor | *Oxtr* | Mm01182684_m1 |
| Purinergic receptor P2X, ligand-gated ion channel, 7 | *P2rx7* | Mm01199500_m1 |
| Prostaglandin-endoperoxide synthase 2 | *Ptgs2* | Mm00478374_m1 |
| Regulator Of G Protein Signaling 4 | *Rgs4* | Mm00501389_m1 |
| Suppressor of cytokine signaling 3 | *Socs3* | Mm00545913_s1 |
| S100 calcium-binding protein A9 | *S100a9* | Mm00656925_m1 |
| Toll-like receptor 4 | *Tlr4* | Mm00445273_m1 |
| Toll-like receptor 9 | *Tlr9* | Mm00446193_m1 |
| Tumor necrosis factor | *Tnf* | Mm00443258_m1 |
| Tumor necrosis factor receptor superfamily member 1a | *Tnfrsf1a* | Mm00441883_g1 |

**Table S3.** Antibodies utilized for flow cytometry

| Antigen/isotype | Fluorochrome | Clone | Company | Catalog number |
|---|---|---|---|---|
| Rat anti-mouse F4/80 | PE-CF594 | T45-2342 | BD Biosciences | 565613 |
| Mouse anti-mouse CD45.1 | FITC | A20 | BD Biosciences | 553775 |
| Mouse anti-mouse CD45.2 | APC-Cy™7 | 104 | BD Biosciences | 560694 |
| Rat IgG2α, κ Isotype Control | PE-CF594 | R35-95 | BD Biosciences | 562302 |
| Mouse IgG2α, κ Isotype Control | FITC | G155-178 | BD Biosciences | 553456 |
| Mouse IgG2α, κ Isotype Control | APC-Cy™7 | G155-178 | BD Biosciences | 557751 |
